# Sod1 Integrates Oxygen Availability to Redox Regulate NADPH Production and the Thiol Redoxome

**DOI:** 10.1101/2021.03.04.433951

**Authors:** Claudia Montllor-Albalate, Hyojung Kim, Alex P. Jonke, Matthew P. Torres, Amit R. Reddi

## Abstract

Cu/Zn superoxide dismutase (Sod1) is a highly conserved and abundant antioxidant enzyme that detoxifies superoxide (O_2_^⸱-^) by catalyzing its conversion to dioxygen (O_2_) and hydrogen peroxide (H_2_O_2_). Using *Saccharomyces cerevisiae* and mammalian cells, we discovered that a major new aspect of the antioxidant function of Sod1 is to integrate O_2_ availability to promote NADPH production. The mechanism involves Sod1-derived H_2_O_2_ oxidatively inactivating the glycolytic enzyme, glyceraldehyde phosphate dehydrogenase (GAPDH), which in turn re-routes carbohydrate flux to the oxidative phase of the pentose phosphate pathway (oxPPP) to generate NADPH. The aerobic oxidation of GAPDH is exclusively dependent on and rate-limited by Sod1. Thus, Sod1 senses O_2_ via O_2_^⸱-^ to balance glycolytic and oxPPP flux, through control of GAPDH activity, for adaptation to life in air. Importantly, this new mechanism for Sod1 antioxidant activity requires the bulk of cellular Sod1, unlike for its role in protection against O_2_^⸱-^ toxicity, which only requires < 1% of total Sod1. Using mass spectrometry, we identified proteome-wide targets of Sod1-dependent redox signaling, including numerous metabolic enzymes. Altogether, Sod1-derived H_2_O_2_ is important for antioxidant defense and a master regulator of metabolism and the thiol redoxome.

**Significance Statement:** Cu/Zn superoxide dismutase (Sod1) is a key antioxidant enzyme and its importance is underscored by the fact that its ablation in cell and animal models results in oxidative stress, metabolic defects, and reductions in cell proliferation, viability, and lifespan. Curiously, Sod1 detoxifies superoxide radicals (O_2_^⸱-^) in a manner that produces an oxidant as a byproduct, hydrogen peroxide (H_2_O_2_). While much is known about the necessity of scavenging O_2_^⸱-^, it is less clear what the physiological roles of Sod1-derived H_2_O_2_ are. Herein, we discovered that Sod1-derived H_2_O_2_ plays a very important role in antioxidant defense by stimulating the production of NADPH, a vital cellular reductant required for ROS scavenging enzymes, as well as redox regulating a large network of enzymes.

## 1. Introduction

Superoxide dismutases (SODs) are a highly conserved class of antioxidant enzymes that serve on the frontline of defense against reactive oxygen species (ROS). SODs, which detoxify O_2_^⸱-^ by catalyzing its disproportionation into O_2_ and H_2_O_2_, are unique amongst antioxidant enzymes in that they also produce a ROS byproduct. While much is known about the necessity of scavenging O_2_^⸱-^, it is less clear what the physiological consequences of SOD-derived H_2_O_2_ are. Paradoxically, increased expression of Cu/Zn SOD (Sod1), which accounts for the majority of SOD activity in cells (1), is actually associated with reduced cellular H_2_O_2_ levels (2), suggesting there may be additional unknown mechanisms underlying Sod1 antioxidant activity.

The cytotoxicity of O_2_^⸱-^ is largely due to its ability to oxidize and inactivate [4Fe-4S] cluster-containing enzymes, which results in defects in metabolic pathways that utilize [4Fe-4S] proteins and Fe toxicity due to its release from damaged Fe/S clusters (3-6). The released Fe can catalyze deleterious redox reactions, and, in particular, production of hydroxyl radicals (^.^OH) via Haber-Weiss and Fenton reactions, which indiscriminately oxidizes lipids, proteins, and nucleic acids (4, 7). The importance of Sod1 in oxidative stress protection is underscored by reduced proliferation, decreased lifespan, and numerous metabolic defects, including cancer, when *SOD1* is deleted in various cell lines and/or organisms (7-11). It was previously proposed that Sod1 limits steady-state H_2_O_2_ levels because of its ability to prevent the O_2_^⸱-^-mediated one-electron oxidation of Fe/S clusters, which results in the concomitant formation of H_2_O_2_ (2, 12, 13). However, since vanishingly small amounts of Sod1, < 1% of total cellular Sod1, is sufficient to protect cells against O_2_^⸱-^ toxicity, including oxidative inactivation of Fe/S enzymes (14-16), any changes in Sod1 expression is not expected to alter H_2_O_2_ arising from O_2_^⸱-^ oxidation of Fe/S clusters. How then can Sod1, an enzyme that catalyzes H_2_O_2_ formation, act to reduce cellular H_2_O_2_ levels?

Two previously reported but unexplained metabolic defects in *sod1*Δ strains of *Saccharomyces cerevisiae* point to a potential role for Sod1 in regulating the production of NADPH, a key cellular reductant required for reductive biosynthesis and the reduction and regeneration of H_2_O_2_ scavenging thiol peroxidases (17) and catalases (18, 19). Yeast strains lacking *SOD1* exhibit increased glucose consumption (20) and defects in the oxidative phase of the pentose phosphate pathway (oxPPP) (21), the primary source of NADPH. Inhibition of key rate-limiting enzymes in glycolysis, including phosphofructose kinase (22), glyceraldehyde phosphate dehydrogenase (GAPDH) (23, 24), and pyruvate kinase (25, 26), reduces glucose uptake (27-29) and increases the concentration of glucose-6-phosphate (G6P), a glycolytic intermediate that is also the substrate for the first enzyme in the oxPPP, G6P dehydrogenase (G6PDH), which in turn increases oxPPP flux and NADPH production (30-35). Taken together, we surmised that Sod1 negatively regulates a rate determining enzyme in glycolysis, thereby accounting for the observed metabolic defects in glucose utilization and the oxPPP in *sod1*Δ cells (20, 21).

GAPDH, which catalyzes a rate-determining step in glycolysis (36, 37), is very abundant (38), and contains a H_2_O_2_-reactive catalytic Cys (*k ∼* 10^2^-10^3^ M^-1^s^-1^), represents a critical redox regulated node that can toggle flux between glycolysis and the oxPPP (32). As such, we hypothesized that a novel aspect of the antioxidant activity of Sod1 is to oxidatively inactivate GAPDH using Sod1-catalyzed H_2_O_2_, which would in turn stimulate NADPH production via the oxPPP and enhance cellular peroxide scavenging by thiol peroxidases. This novel mechanism for Sod1 mediated antioxidant activity would explain a number of prior observations, including the findings that elevated Sod1 expression decreases peroxide levels and loss of *SOD1* increases glucose consumption and attenuates oxPPP activity. In addition, more generally, since Sod1-derived H_2_O_2_ has previously been implicated in the redox regulation of other enzymes, including protein tyrosine phosphatases (39) and casein kinases (15, 16), we also sought to identify proteome-wide redox targets of Sod1.

In the present report, we provide evidence highlighting a new antioxidant function for Sod1-derived H_2_O_2_ in integrating O_2_ availability to control NADPH production to support aerobic growth and metabolism. The mechanism involves the conversion of O_2_ to O_2_^⸱-^ by mitochondrial respiration and an NADPH oxidase, followed by the Sod1-catalyzed conversion of O_2_^⸱-^ to H_2_O_2_, which in turn oxidatively inactivates GAPDH. The inhibition of GAPDH serves to re-route metabolism from glycolysis to the oxPPP in order to maintain sufficient NADPH for metabolism in air. The aerobic oxidation of GAPDH is exclusively dependent on and rate-limited by Sod1, suggesting that it provides a privileged pool of peroxides to inactivate GAPDH under physiological conditions. Lastly, we revealed a larger network of cysteine-containing proteins that are oxidized in a Sod1-dependent manner using mass spectrometry-based redox proteomics approaches. Altogether, these results highlight a new mechanism for O_2_ sensing and adaptation, reveal an important but previously unknown antioxidant role of Sod1 that goes beyond O_2_^⸱-^ scavenging to include the stimulation of aerobic NADPH production, and places Sod1 as a master regulator of proteome-wide thiol oxidation and multiple facets of metabolism.

## 2. Results

### 2.1. Sod1 regulates glycolysis

In many eukaryotes, including *Saccharomyces cerevisiae*, glucose uptake negatively correlates with *p*O_2_ (40-43). Indeed, we find that batch cultures of WT yeast cells grown anaerobically consume more glucose per cell than aerobically grown cultures (**Figures 1A** and **1B**). Media glucose concentration is plotted versus cell density, rather than time, to correct for differences in growth rate (**Figure S1A**). Consistent with previous studies (20), aerobic cultures of *sod1*Δ strains consume more glucose than WT cells (**Figures 1A** and **1B**). However, in the absence of oxygen, both WT and *sod1*Δ cells consume similar amounts of glucose (**Figures 1A** and **1B**).

**Figure 1.**
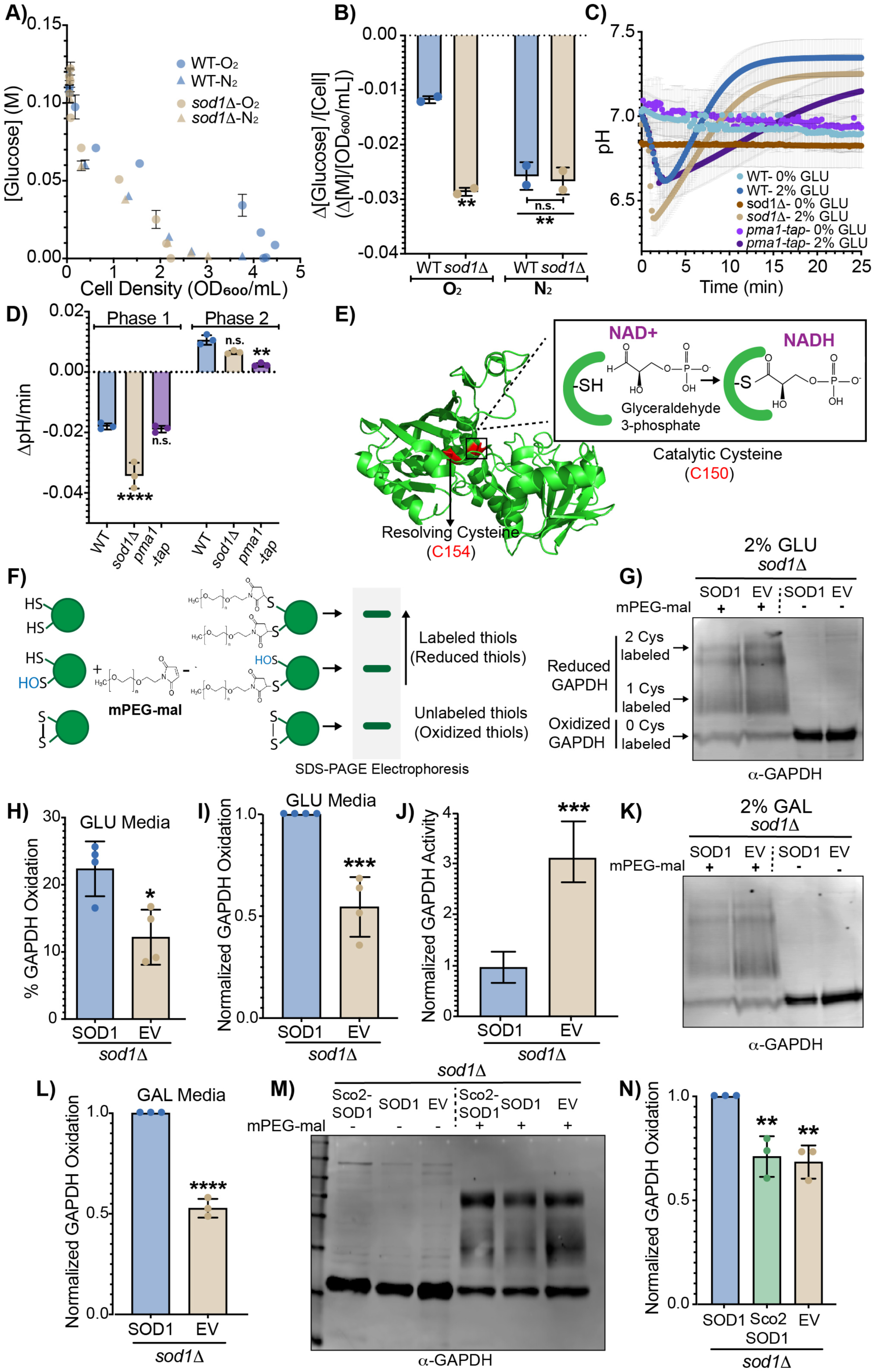
Cytosolic Sod1 regulates glycolysis via the redox regulation of GAPDH. **(A-B)** Extracellular glucose measurements as a function of culture density for WT and *sod1*Δ cells grown aerobically (O_2_) or anaerobically (N_2_) **(A)** and glucose consumption per cell as derived from the slope of the linear regions of the plot **(B)**. The data represent the average ± SD from two biological replicates (See also **Figure S1A**). **(C**-**D**) Time-resolved intracellular pH measurements of glucose starved cells upon pulsing WT, *sod1*Δ, or *pma1-*tap cells expressing a GFP-based pH sensor, pHluorin, with 2% or 0% glucose (GLU) **(C)**. The cytosolic acidification rate, a proxy for glycolytic activity (Phase 1), and the rate of re-alkalization of the cytosol, a proxy for Pma1 activity (Phase 2), is shown for the indicated strains **(D)**. The data represent the average ± SD from triplicate cultures. See also **Figure S1B**. **(E)** The structure of *Saccharomyces cerevisiae* GAPDH, Tdh3 (PDB file 4IQ8), with catalytic Cys_150_ and resolving Cys_154_ represented in red, and a depiction of the Cys_150_ thioester covalent intermediate with glyceraldehyde-3-phosphate. **(F**-**I)** Analysis of Sod1-dependent GAPDH oxidation as assessed by labelling GAPDH with thiol reactive methoxypolyethylene glycol maleimide (mPEG-mal). **(F)** Schematic representation of all possible redox-dependent GAPDH (green spheres) Cys-mPEG-mal adducts and their respective electrophoretic mobilities. **(G)** Representative immunoblot analysis of GAPDH-mPEG-mal adducts in *sod1*Δ cells expressing yeast Sod1 (SOD1) or empty vector (EV) cultured in 2% GLU. **(H)** % GAPDH oxidation, as assessed by quantifying the ratio of mPEG-mal labelled GAPDH to total GAPDH, in the indicated strains from multiple trials. **(I)** Relative GAPDH oxidation in the indicated strains as assessed by normalizing to the % GAPDH oxidation of *SOD1* expressing cells from each trial. Data represents the average ± SD from 4 independent trials. **(J)** Measurements of GAPDH enzymatic activity in *sod1*Δ cells expressing SOD1 or EV. Data represents the average ± SD from quadruplicate cultures. See also **Figures S1O-P**. **(K**-**L)** Assessment of GAPDH oxidation in *sod1*Δ cells expressing yeast Sod1 (SOD1) or empty vector (EV) cultured in 2% galactose (GAL). Representative immunoblot analysis of GAPDH-mPEG-mal adducts **(K)** and normalized GAPDH oxidation from multiple trials **(L)**. Data represents the average ± SD from 3 independent trials. See also Figures S1Q and S1R. **(M**-**N)** Assessment of GAPDH oxidation in *sod1*Δ cells expressing yeast Sod1 (SOD1), mitochondrial IMS targeted Sod1 (Sco2-SOD1), or empty vector (EV). Representative immunoblot analysis of GAPDH-mPEG-mal adducts **(M)** and normalized GAPDH oxidation from multiple trials **(L)** in the indicated strains. Data represents the average ± SD from 3 independent trials. See also **Figure S1S**. The statistical significance is indicated by asterisks using two-tailed Student’s t-tests for pairwise comparisons (panels **H**, **I**, **J**, and **L**) or by one-way ANOVA for multiple comparisons with Dunett’s post-hoc test (panels **B**, **D** and **N**); *p<0.05, **p<0.01, ***p>0.001, ****p<0.0001, n.s.= not significant.

Since defects in glycolytic enzymes, including hexokinase 2 or GAPDH, decrease glucose uptake (27, 28), we sought to determine if the increased glucose consumption of *sod1*Δ cells was associated with altered glycolytic flux. Intracellular cytosolic pH is an excellent reporter of glycolytic activity and can be monitored using the GFP-based ratiometric pH sensor, pHluorin. Glucose starved yeast exhibit an intracellular cytosolic pH of ∼7.0. Upon exposure to glucose, there is a rapid decrease in cytosolic pH to a value of ∼6.6 within 3 minutes, corresponding to proton release in phosphorylation reactions associated with hexokinase, phosphofructose kinase, and GAPDH (**Figures 1C** and **1D**, phase 1) (44). The initial acidification is followed by a slower alkalization phase that brings the cytosolic pH up to ∼7.3 over 15 minutes (**Figures 1C** and **1D**, phase 2), which is due to activation of Pma1, a cell surface H^+^-ATPase that pumps H^+^ into the extracellular space (44). In response to glucose, *sod1*Δ cells exhibit more rapid rates of intracellular acidification, indicating that glycolysis is more active compared to WT cells. Moreover, *sod1*Δ cells exhibit slightly diminished rates of re-alkalization, indicating that PMA1 is less active in response to glucose. Indeed, prior work found that *sod1*Δ cells exhibit a defect in PMA1 activity (16, 45). To rule out that the intracellular acidification phase is unaffected by PMA1-dependent re-alkalization, glucose-dependent changes in intracellular pH were monitored in a strain expressing a hypomorphic allele of PMA1, *pma1-tap*, containing a C-terminal tandem affinity purification (TAP)-tag (**Figure S1B**). In *pma1-tap* cells, the rate of glucose-induced acidification is similar to WT cells (**Figure 1D**, Phase 1), but there is a significant decrease in the rate of re-alkalization (**Figure 1D**, Phase 2). Taken together, the data demonstrate that Sod1 negatively regulates glucose uptake and glycolytic activity.

### 2.2. Extra-mitochondrial Sod1 regulates GAPDH oxidation and activity

We hypothesized that Sod1-derived H_2_O_2_ may negatively regulate glucose uptake and glycolytic activity through the oxidative inactivation of GAPDH, which catalyzes a rate-determining step in glycolysis and contains a peroxide-sensitive active site Cys (23). *Saccharomyces cerevisiae* encodes three GAPDH isoforms, *TDH1*, *TDH2*, and *TDH3*, with *TDH3* being the most highly expressed in log phase cultures, accounting for > 50% of total cellular GAPDH (46). Yeast GAPDH has only two cysteines, catalytic C150 and C154, which sensitizes C150 to oxidation by H_2_O_2_ (**Figure 1E**). In order to probe the Sod1-dependence of GAPDH oxidation, we employed a thiol alkylation assay that exploits the reactivity of methoxypolyethylene glycol maleimide (mPEG-mal) with reduced but not oxidized thiols (47). The extent of GAPDH labeling with mPEG-mal, which is 5 kDa, is assessed by determining the changes in electrophoretic mobility of PEGylated GAPDH, corresponding to single and double labeled GAPDH at reduced C150 and/or C154 (**Figures 1F, 1G** and **S1C**). Thus, the fraction of GAPDH oxidized *in vivo* can be determined by quantifying the ratio of the intensity of unlabeled GAPDH (oxidized GAPDH) to total GAPDH (**Figures S1D** and **S1F-S1M)**. The mPEG-mal approach was validated by treating cells with H_2_O_2_ and observing a 2-fold increase in GAPDH oxidation (**Figure S1D** and **S1E**). Moreover, the identity of the specific sites of mPEG-mal labeling was confirmed by observing that a yeast strain expressing a single allele of Tdh3^C154S^ was found to have only two GAPDH proteoforms corresponding to single and unlabeled GAPDH (**Figure S1N**).

To determine if Sod1 oxidizes GAPDH *in vivo*, we analyzed GAPDH PEGylation in *sod1*Δ cells expressing empty vector (EV) (*sod1*Δ + EV) or WT *SOD1* (*sod1*Δ + *SOD1*). Interestingly, *sod1*Δ + EV cells exhibit an increase in mPEG-mal labeling compared to *sod1*Δ + *SOD1* cells (**Figures 1G-1I**), indicating that GAPDH is more oxidized in cells expressing *SOD1*. Over 4-independent trials, we found that cells expressing *SOD1* exhibit a ∼2-fold increase in GAPDH oxidation compared to cells not expressing *SOD1* (**Figure 1H**). Due to variations in the absolute levels of GAPDH oxidation across multiple trials (**Figure 1H**), we also chose to normalize oxidation levels to *SOD1* expressing cells within each trial (**Figure 1I**). Such an analysis accounts for trial-to-trial variation in GAPDH oxidation and increases the statistical significance of the results between various strains.

We next evaluated the effect of Sod1-mediated GAPDH oxidation on GAPDH catalytic activity. GAPDH catalyzes the oxidative phosphorylation of GAP to 1,3 BPG and requires Cys^150^ for activity. GAPDH activity is 3-fold greater in cells lacking *SOD1* (*sod1*Δ + EV) (**Figures 1J, S1O** and **S1P**), which correlates with the 2-fold decrease in GAPDH oxidation relative to *SOD1* expressing cells (**Figures 1G** and **1I**).

Notably, Sod1-mediated GAPDH oxidation is not dependent on the carbon source. In yeast, glucose represses mitochondrial respiration and promotes fermentation. Galactose is a fermentable carbon source that alleviates respiration repression, resulting in more mitochondrial respiratory activity (16). The absolute (**Figure S1R**) and relative (**Figure 1L**) amounts of GAPDH oxidation in *sod1*Δ + EV and *sod1*Δ + *SOD1* cells cultured in 2% galactose is similar to cells cultured in 2% glucose. (**Figures 1K, 1L, S1Q** and **S1R)**.

We next evaluated the effect of Sod1 localization on GAPDH oxidation. Sod1 is primarily cytosolic but is also present in the mitochondrial IMS. *sod1*Δ cells expressing an IMS-targeted allele of Sod1, Sco2-SOD1 (15, 16, 48), exhibit comparable GAPDH oxidation to *sod1*Δ + EV cells, both of which are significantly lower than cells expressing WT *SOD1* (**Figures 1M, 1N** and **S1S**). Altogether, these results indicate that extramitochondrial Sod1-derived H_2_O_2_ oxidizes GAPDH and decreases its catalytic activity, thereby explaining the previous observations that *sod1*Δ cells exhibit increased glucose uptake (**Figures 1A** and **1B**) and glycolytic activity (**Figures 1C** and **1D**).

### 2.3. Yno1 and mitochondrial respiration are sources of superoxide for GAPDH oxidation

Sod1 requires a superoxide source to catalyze peroxide production for the control of GAPDH oxidation and activity. As with higher eukaryotes, two primary sources of superoxide in yeast include mitochondrial respiration and the yeast NADPH oxidase, Yno1 (49). Both sources contribute towards GAPDH oxidation as respiration deficient *rho^0^* and *yno1*Δ cells exhibit a ∼3-fold lower degree of GAPDH oxidation relative to WT cells and phenocopy the *sod1*Δ mutant (**Figures 2A**, **2B** and **S2A**). Furthermore, overexpression of Yno1 on a galactose-inducible promoter (pYES-YNO1, 3%GAL), which resulted in a ∼30% increase in DHE-detectable superoxide (**Figure S2B**), promoted GAPDH oxidation by 2-fold as compared to cells expressing empty vector (pYES2-EV, 0 and 3%GAL) or that were cultured in non-inducing media (pYES2-YNO1, 0%GAL) (**Figures 2C**, **2D** and **S2C**). In total, these results indicate that both Yno1 and mitochondrial respiration are sources of the superoxide substrate required by Sod1 to drive the H_2_O_2_ dependent oxidation of GAPDH.

**Figure 2.**
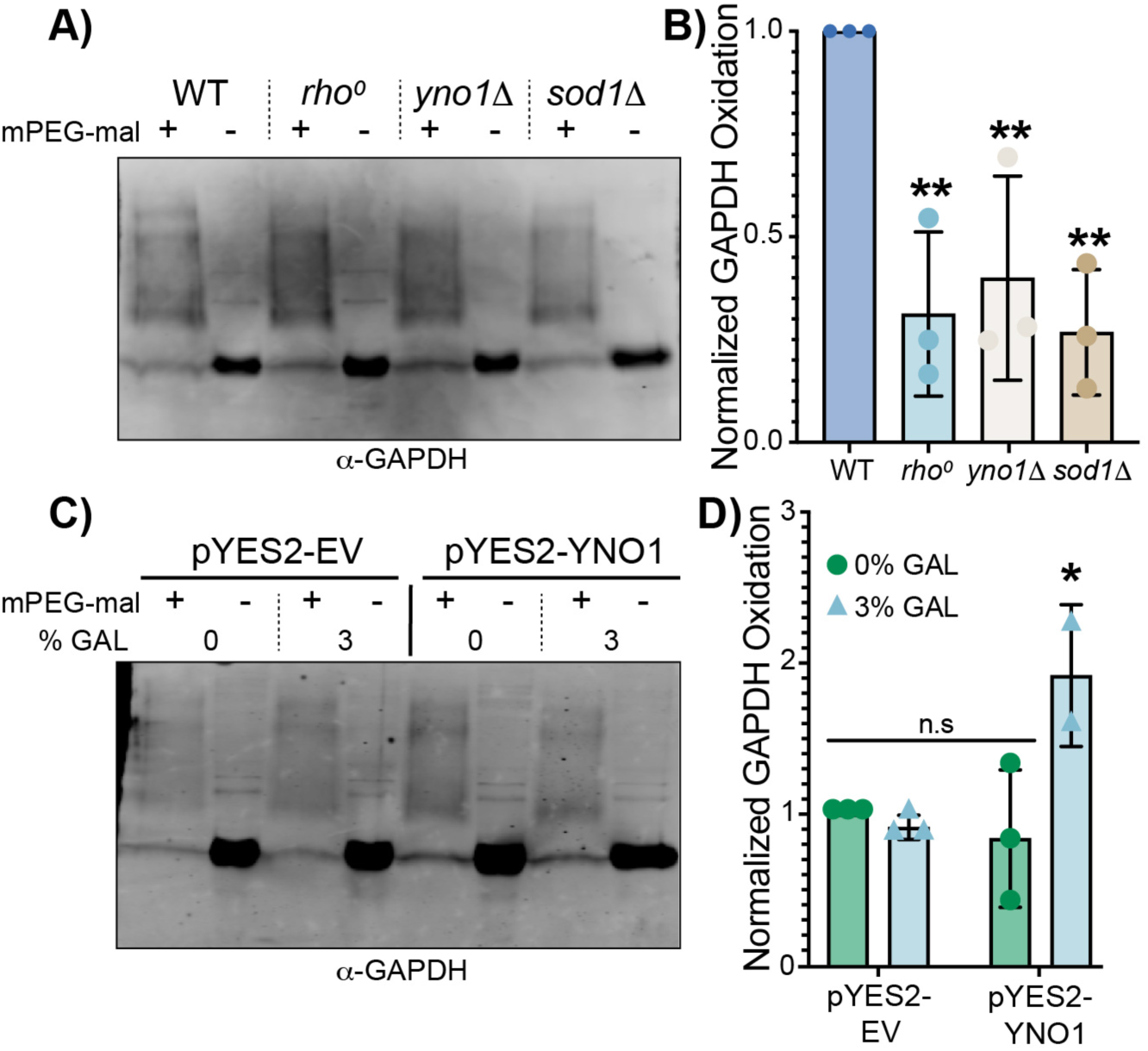
Yno1 and mitochondrial respiration are sources of superoxide for GAPDH oxidation. **(A-B)** Representative immunoblot analysis of GAPDH oxidation as assessed by mPEG-mal labeling of GAPDH in WT, *rho^0^*, *yno1*Δ and *sod1*Δ cells cultured in 2% GLU **(A)** and normalized GAPDH oxidation from multiple trials **(B)**. Data represents the average ± SD from 3 independent trials. See also **Figure S2A**. **(C-D)** Representative immunoblot analysis of GAPDH oxidation as assessed by mPEG-mal labeling of GAPDH in WT cells expressing GAL1-driven *YNO1* (pYES2-YNO1) or empty vector (pYES2-EV) cultured in 2% raffinose, supplemented with non-inducing (0%) or inducing (3%) GAL concentrations and **(C)** the normalized GAPDH oxidation from multiple trials **(D)**. Data represents the average ± SD from 3 independent cultures. See also **Figures S2B-S2C**. The statistical significance relative to WT (panel **B**) or pYES2-EV 0%GAL (panel **D**) is indicated by asterisks using ordinary one-way ANOVA or two-way ANOVA with Dunett’s post-hoc test for the indicated pairwise comparisons in panel **B** and **D**, respectively; *p<0.05, **p<0.01, ***p>0.001, ****p<0.0001, n.s.= not significant.

### 2.4. O_2_ dependent GAPDH oxidation is exclusively dependent on and rate limited by SOD1

All metabolic sources of superoxide and peroxide are ultimately derived from O_2_. We therefore sought to determine if Sod1 is the sole enzymatic adapter that links oxygen availability to the control of GAPDH oxidation and if GAPDH oxidation is rate-limited by Sod1. Towards this end, we first asked if Sod1 mediates O_2_-dependent GAPDH oxidation. Since WT Sod1 is transcriptionally and post-translationally down-regulated in response to hypoxia and anoxia (50-53), we utilized *sod1*Δ cells expressing *ADH1*-driven Sod1^P144S^, which is a mutant previously engineered to constitutively express mature enzymatically active Sod1 even in the absence of O_2_ (16, 52). Indeed, the Sod1^P144S^ mutant is enzymatically active in lysates derived from both aerobic and anaerobic cultures, whereas WT Sod1 is only fully active in lysates derived from aerobically cultured cells (**Figures 3A** and **3B**). *sod1*Δ cells expressing WT or Sod1^P144S^ exhibit a nearly 2-fold decrease in GAPDH oxidation when cultured anaerobically, consistent with the requirement for O_2_ as the metabolic origin of superoxide and peroxide (**Figures 3C**, **3D** and **S2D**). However, remarkably, the O_2_-dependence of GAPDH oxidation is completely lost in cells lacking *SOD1*. Furthermore, we determined that the oxidation of GAPDH is rate limited by Sod1, finding that GAL-regulated titration of Sod1 levels results in a dose-dependent increase in GAPDH oxidation (**Figures 3E, 3F** and **S2E**). Altogether, these results indicate that the aerobic oxidation of GAPDH is exclusively dependent on and rate limited by Sod1.

**Figure 3.**
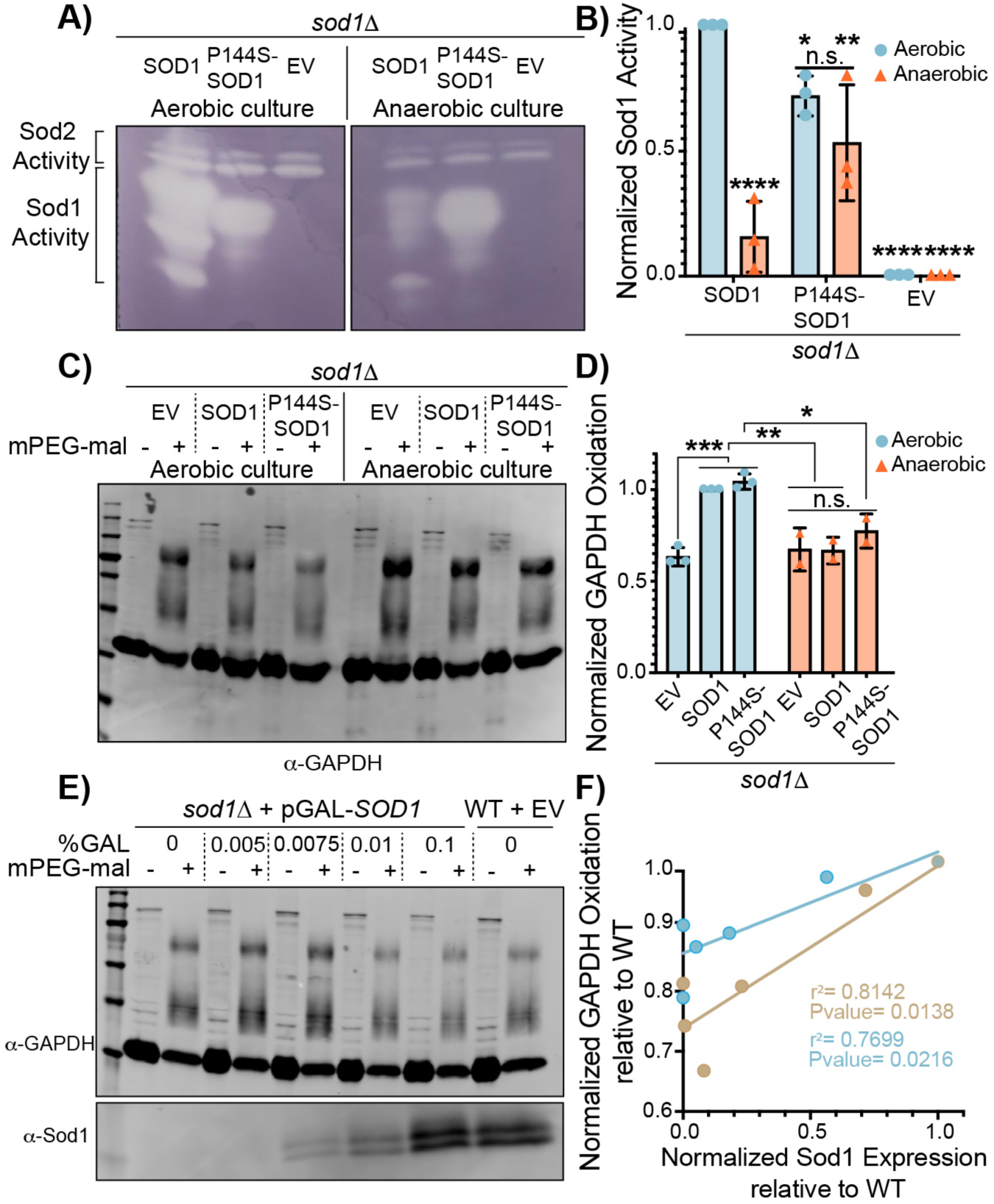
O_2_ dependent GAPDH oxidation is exclusively dependent on and rate limited by Sod1. **(A-D)** Assessment of the Sod1-dependence on the aerobic oxidation of GAPDH. **(A)** Representative SOD activity gel to assess aerobic and anaerobic Sod1 maturation in *sod1*Δ cells expressing empty vector, WT SOD1, or the P144S *sod1* mutant and **(B)** normalized levels of mature Sod1 from multiple trials. Data represents the average ± SD from 3 independent cultures. **(C)** Representative immunoblot analysis of GAPDH oxidation as assessed by mPEG-mal labeling of GAPDH in aerobic or anaerobic *sod1*Δ cells expressing empty vector, WT SOD1, or the P144S *sod1* mutant and **(D)** the normalized GAPDH oxidation from multiple trials. Data represents the average ± SD from two or three independent trials. See also **Figure S2D**. **(E-F)** Representative immunoblot analysis of GAPDH oxidation in WT or *sod1*Δ cells expressing GAL-driven *SOD1* cultured with increasing concentrations of galactose (GAL) (0%, 0.005%, 0.0075%, 0.01% and 0.1% GAL) and **(E)** the positive correlation between Sod1 expression and GAPDH oxidation from two independent trials **(F)**. Sod1 expression and GAPDH oxidation were normalized to that of the WT cells and the linear regression analysis of the two trials gives coefficients of determination (r^2^) of .81 and .77, with p-values of .01 and .02, respectively. See also **Figure S2E**. The statistical significance relative to the aerobic SOD1 expressing cells is indicated by asterisks using 2-way ANOVA for multiple comparisons with Dunett’s or Bonferroni post-hoc test for the indicated pairwise comparisons in panel **B** and **D**, respectively; *p<0.05, **p<0.01, ***p>0.001, ****p<0.0001, n.s.= not significant.

### 2.5. Sod1-mediated oxidative inactivation of GAPDH results in increased NADPH production and oxidative stress resistance

We next determined if the Sod1-dependent oxidative inactivation of GAPDH results in re-routing of glycolytic metabolism towards oxPPP to produce NADPH and increase resistance to oxidative stress. Titration of Sod1 expression using a GAL-driven *SOD1* allele results in both a dose dependent increase in GAPDH oxidation as well as NADPH levels (**Figures 4A-4C** and **S3A-S3F**). Control experiments in which GAL is titrated into cells expressing a non-GAL-driven *SOD1* allele indicate that GAL alone does not alter NADPH levels (**Figure S3G**). Moreover, *tdh3*Δ cells, which express ∼60% less GAPDH than WT cells, exhibit increased NADPH levels, consistent with our finding that oxidative inactivation of GAPDH increases NADPH (**Figures 4D-4F**).

**Figure 4.**
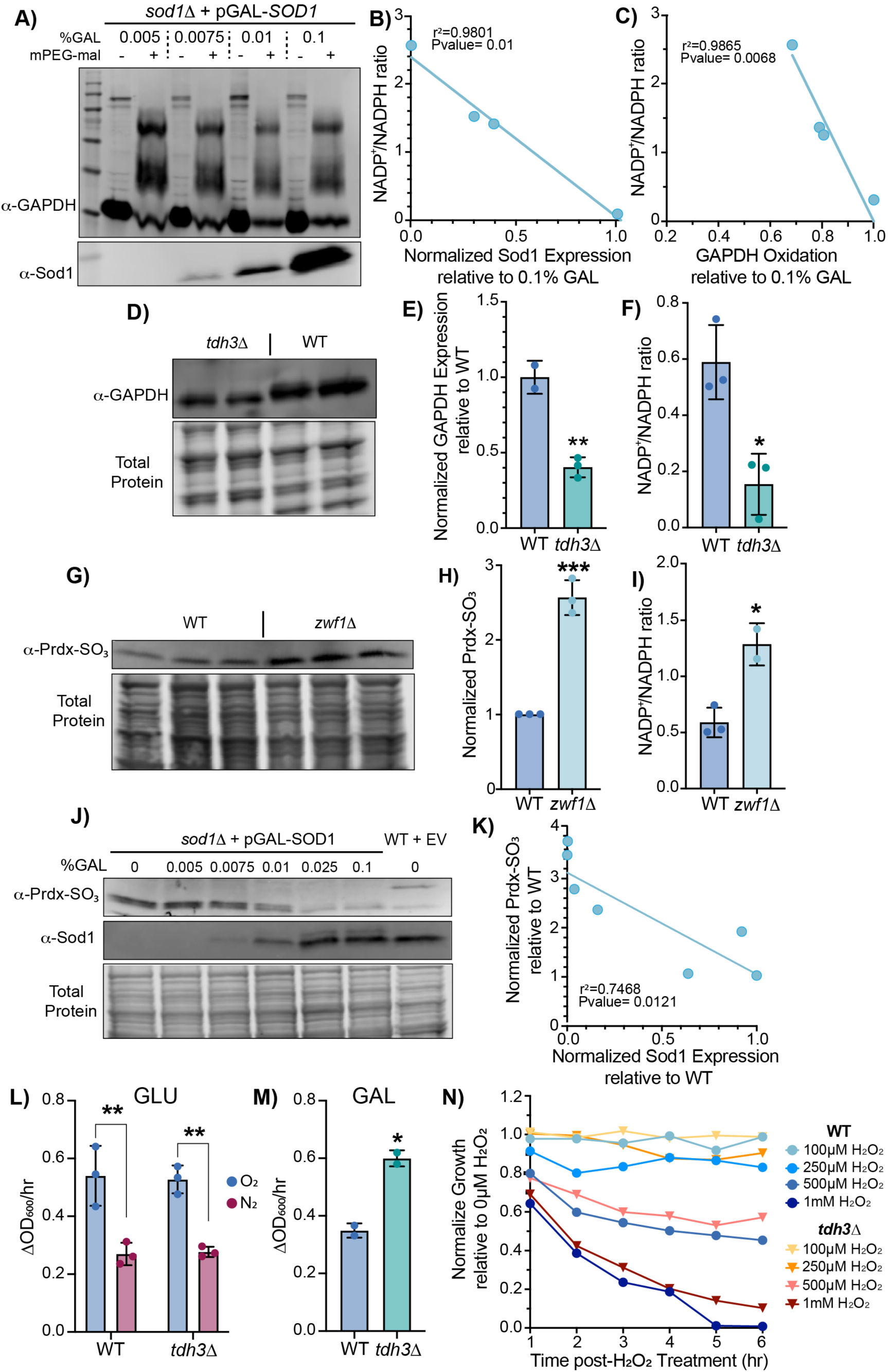
Sod1-mediated oxidative inactivation of GAPDH results in increased NADPH production and resistance to oxidative stress. **(A-C)** Sod1 expression positively correlates with GAPDH oxidation and NADPH production. **(A)** Immunoblot analysis of GAPDH oxidation as assessed by mPEG-mal labeling of GAPDH in *sod1*Δ cells expressing GAL-driven *SOD1* cultured with increasing concentrations of galactose (GAL) (0.005%, 0.0075%, 0.01% and 0.1% GAL). See also **Figure S3A** (**B**-**C**) The NADP+/NADPH ratio inversely correlates with normalized Sod1 expression **(B)** and normalized GAPDH oxidation **(C)**. The linear regression analysis in panels **B** and **C** gives coefficient of determinations (r^2^) of .98 and .99 and p-values of .01 and .007, respectively. See also **Figures S3B-S3H**. **(D-F)** Depletion of intracellular GAPDH decreases the NADP+/NADPH ratio. **(D)** Representative immunoblot analysis of GAPDH expression in WT and *tdh3*Δ cells and **(E)** normalized GAPDH expression from multiple trials. **(F)** Measurements of the NADP+/NADPH ratio in WT and *tdh3*Δ. Data represents the average ± SD from triplicate cultures. **(G-I)** Ablation of glucose-6-phosphate dehydrogenase (Zwf1) increases peroxiredoxin oxidation (Prdx-SO_3_) and the NADP/NADPH ratio. Representative immunoblot analysis of Prdx-SO_3_ **(G)**, normalized Prdx-SO_3_ levels from replicates, and NADP+/NADPH ratios from WT and *zwf1*Δ cells. Data represents the average ± SD from triplicate cultures. **(J-K)** Sod1 expression inversely correlates with Prdx-SO_3_. **(J)** Representative immunoblot analysis of peroxiredoxin oxidation (Prdx-SO_3_) and Sod1 expression in WT or *sod1*Δ cells expressing GAL-driven *SOD1* cultured with increasing concentrations of galactose (GAL) (0%, 0.005%, 0.0075%, 0.01%, 0.025%, and 0.1% GAL). **(K)** Sod1 expression inversely correlates with Prdx oxidation, with a linear regression analysis giving a coefficient of determination (r^2^) of 0.75 and a p-value of .01. **(L-N)** Aerobic and anerobic growth rates in 2%Glucose media **(L)**, aerobic growth rate in 2%GAL media **(M)** and peroxide sensitivity of WT and *tdh3*Δ cells **(N)**. Data represent the average ± SD from triplicate cultures. The statistical significance relative to WT is indicated by asterisks using two-tailed Student’s t-test for pairwise comparisons in panels **E**, **F**, **H**, and **I**; *p<0.05, ***p>0.001.

Since NADPH is required for the reduction and regeneration of numerous cellular antioxidant systems, we also determined if the alterations in Sod1-mediated NADPH production correlated with the oxidation state of Tsa1, a yeast peroxiredoxin important for oxidant defense and redox signaling that is maintained in a reduced state using reducing equivalents from NADPH. Consistent with a role for NADPH in maintaining reduced Tsa1, *zwf1*Δ cells lacking glucose-6-phosphate dehydrogenase (G6PD), which catalyzes the first committed step of the oxPPP, exhibit decreased NADPH levels and elevated Tsa1 oxidation as assessed by immunoblotting using an antibody that recognizes sulfinic acid oxidized peroxiredoxins (Prx-SO_3_) (**Figure 4G-4I**). Titration of Sod1 using a GAL-regulated promoter results in a dose dependent decrease in Tsa1 oxidation, consistent with the role of Sod1 in promoting NADPH production due to oxidative inactivation of GAPDH (**Figure 4J** and **4K**). In order to determine if the oxidative inactivation of GAPDH by Sod1 provides a physiological benefit for cells, we measured the aerobic growth and oxidative stress resistance of cells expressing low and high levels of GAPDH. Although WT and *tdh3*Δ cells have similar aerobic and anaerobic growth rates in 2% GLU (**Figure 4L**), when cultured in 2% GAL, a fermentable carbon source that promotes respiration in yeast, *tdh3*Δ cells have a marked enhancement in growth rate compared to WT cells (**Fig. 4M**). Moreover, *tdh3*Δ cells exhibit greater resistance to peroxide stress (**Figure 4N**). Altogether, the enhanced aerobic fitness and peroxide tolerance of cells depleted of GAPDH is consistent with a beneficial role for the Sod1-mediated inactivation of GAPDH.

### 2.6. SOD1 regulates GAPDH oxidation in human cells

Both Sod1 and GAPDH are highly conserved from yeast to humans. To establish the conservation of the Sod1-GAPDH redox-signaling axis, we employed human embryonic kidney HEK293 cells. It is worth noting that human GAPDH has an additional peroxide reactive Cys, which would result in 3 PEGylated proteoforms, corresponding to triple-, double-, and single-labeled GAPDH in immunoblots (**Figures S4A-S4C**). As expected, cells treated with H_2_O_2_ exhibited less mPEG-mal labelling, indicating a larger fraction of oxidized GAPDH compared to non-treated cells (**Figure S4B**). To determine if Sod1 promoted GAPDH oxidation, we depleted Sod1 in HEK293 cells using small interfering RNA against Sod1 (siSOD1) or scrambled control RNAi (siCTRL). Across various trials, we consistently observed a depletion of ∼60-80% of Sod1 (**Figures 5A** and **5B)** and a corresponding decrease in GAPDH oxidation (**Figures 5C, 5D, S4C** and **S4D**). The extent to which Sod1 promotes GAPDH oxidation ranges from ∼7 to 16%, which is comparable to the contribution observed from exogenous peroxide treatment (**Figure S4B**). Moreover, as with yeast, we find that Sod1 expression levels across multiple trials positively correlate with GAPDH oxidation (**Figure 5E**). The correlation coefficient (r^2^) from a linear regression analysis is 0.7605, with p = .0022. Notably, the y-intercept of the linear regression is close to 0, indicating that in the complete absence of Sod1, GAPDH oxidation is expected to be ∼0% (**Figure 5E**). Furthermore, cell lines that overexpress Sod1, such as the breast cancer cell line MCF7 (54), exhibit a nearly 3-fold increase in GAPDH oxidation compared to HEK293 cells (**Figure S4E** and **S4F**). Altogether, these results indicate that Sod1-mediated oxidation of GAPDH is conserved in humans.

**Figure 5.**
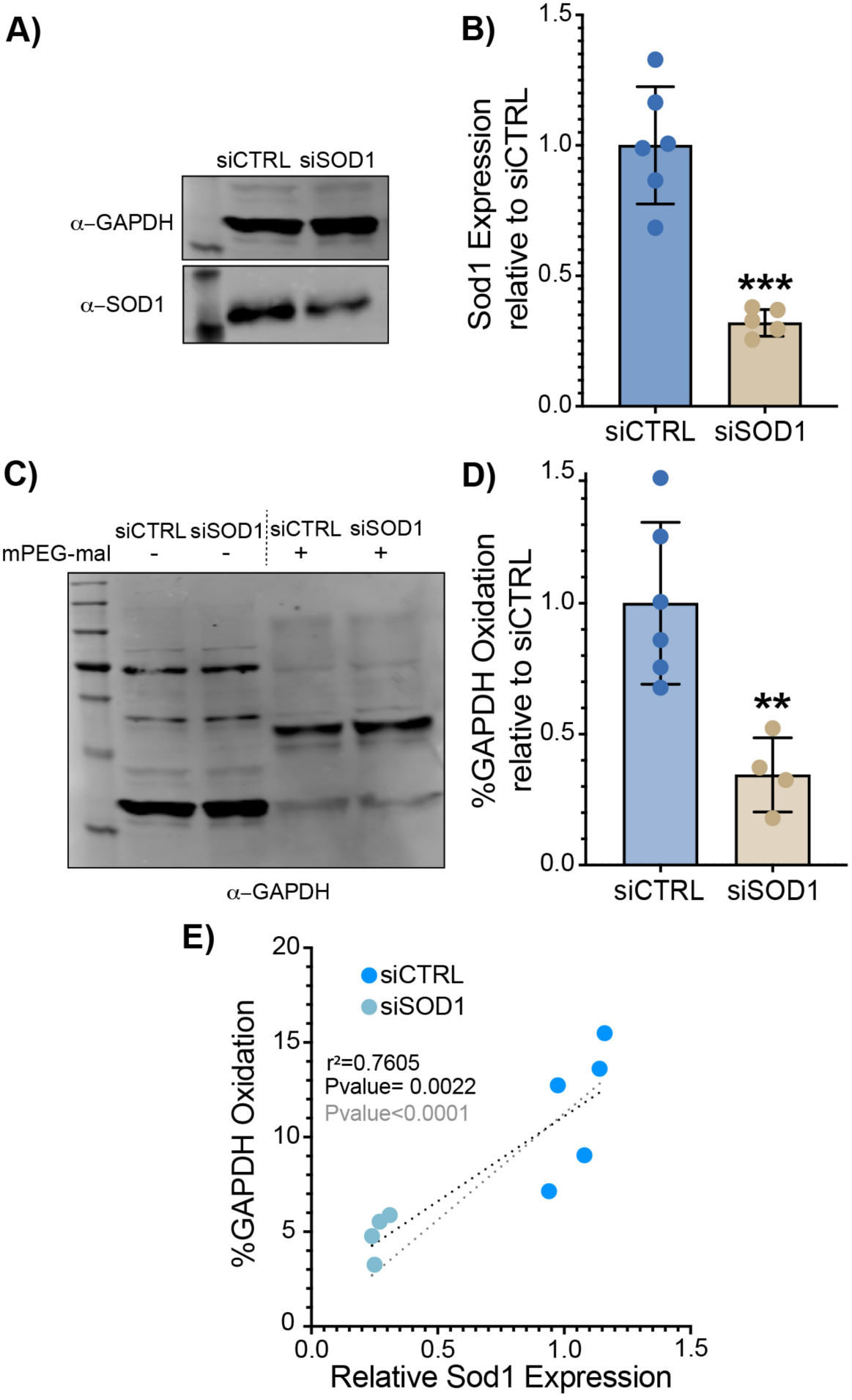
Sod1-mediated GAPDH oxidation is conserved in human cells. **(A-B)** Representative immunoblot analysis of Sod1 and GAPDH from human embryonic kidney HEK293 cells that have *SOD1* silenced with small interfering RNA_i_ (siSOD1) or control scrambled RNA_i_ (siCTRL) (**A)** and normalized ratios of Sod1/GAPDH expression from multiple trials **(B)**. Data represents individual values from five or six biological replicates. **(C**-**D)** Representative immunoblot analysis of GAPDH oxidation as assessed by mPEG-mal labeling of HEK293 cells with silenced (siSOD1) or unsilenced (siCTRL) Sod1 **(C)** and normalized GAPDH oxidation from multiple trials **(D)**. Data represents the average ± SD from five or six biological replicates. See also **Figures S4C-S4D**. **(E)** The correlation between %GAPDH oxidation and relative Sod1 expression, as assessed by measuring the ratio of Sod1/GAPDH levels, from the multiple trials depicted in panel **D**. A linear regression analysis gives a coefficient of determination (r^2^) of .76 and a p-value of 0.002 (black line). If the x and y-intercept of the linear regression is fixed at 0% oxidation and no Sod1 expression (gray line), the correlation remains significant (p<0.0001). In panels **B** and **D**, the statistical significance relative to siCTRL is indicated by asterisks using two-tailed Student’s t-test for pairwise comparison; *p<0.05, **p<0.01, ***p>0.001, ****p<0.0001, n.s.= not significant.

### 2.7. Redox proteomics identifies additional putative targets of Sod1 redox regulation

The high abundance and broad cellular distribution of Sod1 suggests that it may function as a redox regulator of a broad variety of substrates in addition to GAPDH. To test this, we conducted a high-powered quantitative redox proteomics screen of WT and *sod1*Δ yeast to identify Sod1-dependent redox substrates. We used a combined SILAC-TMT approach, whereby *sod1*Δ-dependent changes in protein abundance are quantified through Stable Isotope Labeling with Amino acids in Cell culture (SILAC) and changes in reversible cysteine oxidation are quantified through cysteine-reactive Iodoacetyl Tandem Mass Tags (iodo-TMT) before and after reduction with DTT (**Figure 6A**). In this case, the same peptide from WT and *sod1*Δ cells is distinguished by a mass shift allowing quantitative comparison of the peptide’s abundance in each strain (Protein expression difference = log_2_ (*sod1*Δ_light_/WT_heavy_). Subsequent fragmentation/sequencing of each peptide reveals the peptide/protein identity and, if cysteine is present, releases TMT reporter ions enabling quantification of the cysteine oxidation percentage in each strain (Oxidation difference = *sod1*Δ%Ox -WT%Ox; *see methods*). Consequently, the study reveals a broad range of proteins that undergo significant changes in abundance, cysteine oxidation, or both (**Figure 6B** and **S5A**).

**Figure 6.**
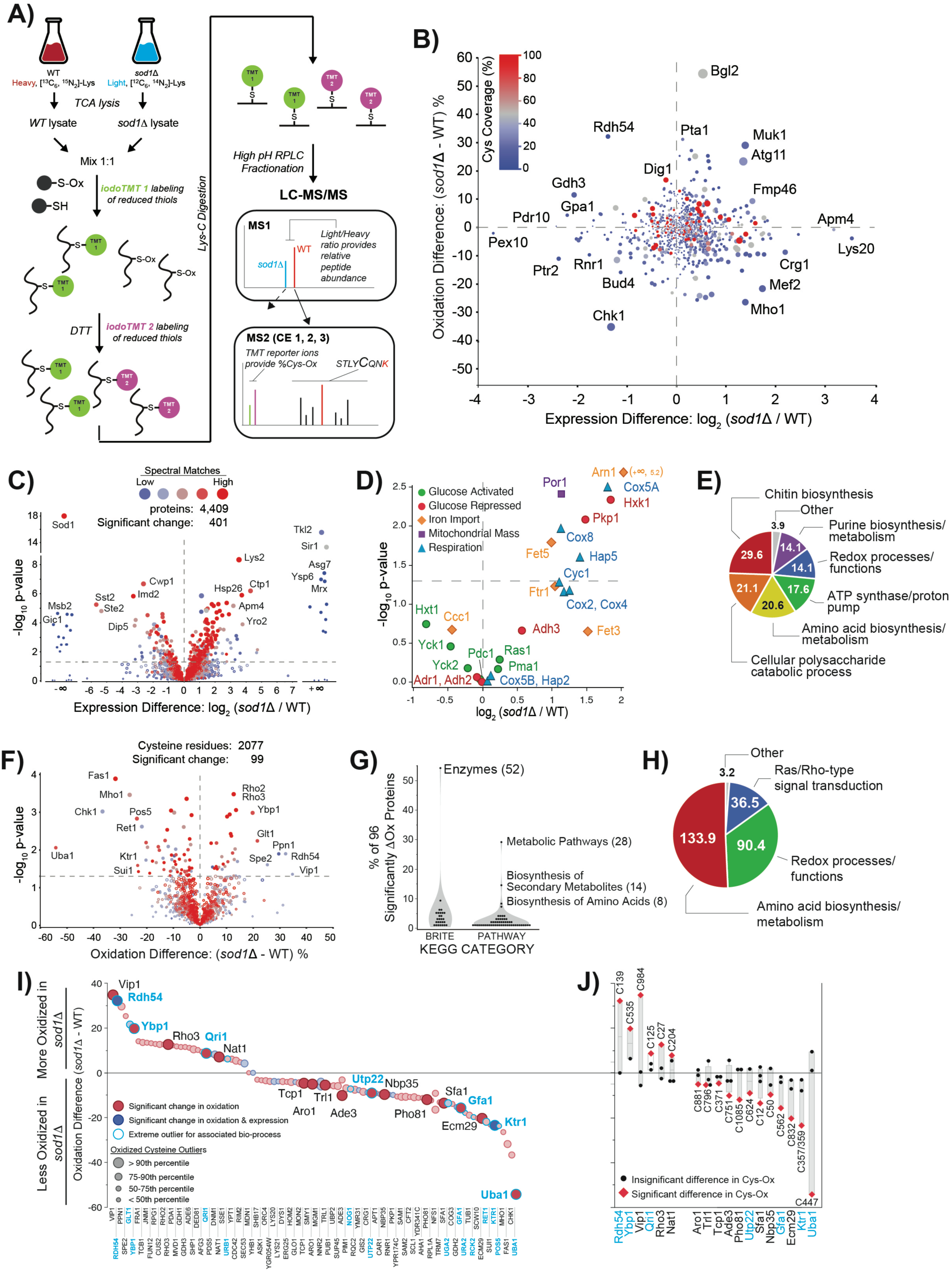
Redox proteomics identifies additional putative targets of Sod1 redox regulation. **(A)** SILAC-TMT redox proteomics workflow. SILAC labels distinguish the cellular origin of each peptide. Cysteine-reactive and isobaric iodoTMT reporters, undetectable during the first MS stage (MS1) and released during peptide fragmentation in MS2, enable unique cysteine %oxidation calculation in WT and *sod1*Δ cells. **(B)** Plot of aggregated average cysteine oxidation difference relative to protein expression difference in *sod1*Δ versus WT cells. Points near the origin (0,0) undergo less change in relative abundance and cysteine oxidation compared to points further from the origin. Cysteine coverage indicates the fraction of cysteine residues detected for each protein. **(C)** Volcano plot of -Log_10_(P-value) relative to protein expression difference [Log_2_(*sod1*Δ/WT)] calculated from aggregate SILAC data. Proteins above -Log_10_(P-value) 1.3 are significantly different in abundance between the two strains (p<0.05). Positive and negative values of log_2_(*sod1*Δ/WT) indicate proteins that are more expressed in *sod1*Δ and WT, respectively. Proteins with a single detected label are indicated as +/-infinity. **(D)** Subset of data from C, showing expression change for proteins associated with bioprocesses functionally associated with loss of *SOD1*. See also **Figure S5B (E)** Median fold enrichment values for gene ontology (GO) bins observed for proteins undergoing >2-fold change in abundance between *sod1*Δ and WT cells. **(F)** Volcano plot of -Log_10_(P-value) relative to protein expression difference [%Ox*_sod1_*_D_ -%Ox_WT_] calculated from iodoTMT data aggregated at the cysteine level. -Log_10_(P-value) of 1.3 or above indicate significantly different Cysteine oxidation between the two strains (p<0.05). See also **Figure S6A (G)** KEGG Brite and Pathway enrichment analysis for the 96 proteins harboring oxidation sites that change significantly in the absence of *SOD1*. Number of associated proteins shown in parentheses. See also **Figure S5D (H)** Median fold enrichment values for GO bins observed for proteins with at least one cysteine undergoing >5% difference in oxidation between WT and *sod1*Δ. **(I)** Rank-ordered plot of cysteine residues from proteins that exhibit statistically significant differences in oxidation between *sod1*Δ and WT. Statistical outliers in oxidation change between strains (circle size) and with respect to other cysteine residues contained within share bioprocess (blue halo, blue text) are indicated. **(J)** Deconvoluted oxidation differences for proteins in which a single cysteine was found to be a statistical outlier from other observed cysteine oxidation sites in the same protein. See also **Figure S6B**.

Focusing first on the effects of SOD1 deletion on proteome-wide protein abundance, we independently analyzed the SILAC MS data. A total of 4,409 proteins were confidently detected and quantified, and ∼9% of these (373) exhibited significant differences in abundance between the two strains (**Figure 6C, Table S1A**). Of these, 114 (30.6%) exhibit a significant decrease in protein abundance in *sod1*Δ cells compared to 259 (69.4%) proteins that exhibit a significant increase in abundance. The changes in protein expression reflect known metabolic defects associated with loss of *SOD1*, including alleviation of glucose repression, increased mitochondrial mass, induction of the iron starvation and antioxidant responses, and diminished plasma membrane casein kinase expression **(Figure 6D** and **S5B)**. GO enrichment analysis of proteins undergoing a statistically significant change in abundance revealed a set of outlier ontologies (i.e. GO Terms falling outside the 90^th^ percentile in the distribution of fold enrichment) that could be clustered into discrete bins. Notably, we observed the greatest enrichment for ontologies linked to glycolysis and pentose phosphate pathways. This included chitin synthesis (Median Fold Enrichment (FE): 29.6), which feeds directly from D-fructose-6-phosphate – a precursor to the GAPDH substrate D-glyceraldehyde-3-phosphate and a key metabolite in the PPP; cellular polysaccharide catabolism (starch and sucrose catabolism) (FE: 21.1); amino acid biosynthesis (FE: 20.6), which relies on several glycolytic intermediates; as well as purine biosynthesis/metabolism that feeds on D-ribose-5-phosphate from the PPP (FE: 14.1) (**Figure 6E, Tables S1B** and **S1C**). We observed lower but still significant GO enrichment of proteins involved in redox processes and functions (FE: 14.1), including ontologies encompassing oxidoreductase activity, NADH oxidation, and ROS metabolic processes; and enrichment of proteins associated with ATP synthase/proton pumping (FE: 17.6).

Next, we focused on the effects of *SOD1* deletion on proteome-wide cysteine oxidation as revealed by iodoTMT MS data. A total of 2,077 cysteine residues were confidently detected ranging in oxidation level from 1 to 100% and a median level of 28.9% across all sites measured. Cross-referencing all sites against protein functional residues revealed 54 of these sites associated with metal binding (38, 74%), active sites (11, 20%), and protein or DNA binding (5, 9%) and cysteine residues involved in these functions also exhibited statistically higher overall oxidation levels in both WT and *sod1*Δ (44.6%, p<0.0001) (**Figure S5C** and **Table S1D**). Global oxidation levels were nearly identical between both strains **(Figure S5B**).

Approximately 5% of the quantified cysteine sites (99 sites, 96 proteins) undergo significant changes in oxidation between WT and *sod1*Δ cells, and of these, we found that 65% exhibit reduced oxidation levels in the absence of *SOD1* (**Figure 6F** and **Table S1E**). GAPDH peptides, and C150/C154 specifically, were readily detected but failed to undergo iodoTMT reporter release under any of the three collision energies used for peptide fragmentation, preventing quantitation of SOD1-dependent oxidation in these experiments. On average, cysteine residues in *sod1*Δ cells appear to be less oxidized compared to WT cells, consistent with a role for Sod1 in providing a source of peroxides for thiol oxidation. On the other hand, cysteine residues that are more oxidized in *sod1*Δ cells compared to WT may be due to oxidative stress, in part mediated by the decrease in NADPH production due to diminished Sod1-mediated GAPDH oxidation and the subsequent reduction in flux through the oxPPP.

Over half of the proteins harboring the 99 significantly changing oxidation sites are classified as enzymes, with the next most populous class represented at less than 10% in comparison (**Figure 6G, Tables S1F** and **S1G**). Most of the 96 proteins are also associated with metabolism, synthesis of secondary metabolites, and biosynthesis of amino acids. Clustering of ontologies that were statistical outliers for fold enrichment supported these findings and produced 3 distinct ontology bins. Ontologies associated with amino acid biosynthesis were predominantly enriched (FE: 133.9) and to a far greater extent than what we observed for Sod1-dependent protein abundance changes (**Figure 6H, Tables S1C** and **S1H**). Moreover, many of the proteins within this bin exhibit reduced oxidation in the absence of *SOD1*, and map to multiple different amino acid biosynthesis/metabolism pathways including those for lysine (Lys20/21), arginine (Car1), threonine (Hom2), and s-adenosyl methionine (Sam1/2) to name a few (**Figure S5D** and **S5E**). Ontologies encompassing redox processes and functions were also highly enriched (FE: 90.4), and include proteins involved in catalysis of redox reactions in which a CH-NH2 group acts as a hydrogen or electron donor and reduces NAD+ or NADP+. Lastly, we also found significant enrichment of proteins involved in Ras and Rho signal transduction (median FE: 36.5), which hint at a unique and unrealized connection between *SOD1*, redox homeostasis, actin mobilization, and cell motility.

Considering the exclusive role of Sod1 as a direct regulator of GAPDH through C150 oxidation, we asked whether our proteomic data might contain evidence of a similar relationship between Sod1 and other proteins, wherein: one, the protein is specifically affected at one out of multiple possible cysteine residues; and two, oxidation of this cysteine is significantly reduced in the absence of *SOD1*. To create this filter, we binned each of the 99 sites based on whether the change in oxidation was a statistical outlier (>90^th^ percentile) from other cysteine residues in the protein, revealing 18 proteins (**Figure 6J**). In 7 of these cases, the change in oxidation was also found to be an outlier with respect to the associated bioprocess for the given protein (**Figure 6I** blue halos, **Figure S6A** and **Table S1I**). Deconvolution of their site-specific changes in oxidation revealed a range of responses reflecting the sensitivity of specific cysteine residues to the presence or absence of *SOD1* (**Figure 6J**). Some of these coincided with proteins enriched previously through ontology, most notably Sfa1 and Gfa1 involved in pyruvate metabolism and conversion of D-fructose-6-phosphate, respectively. Sfa1-C12 and Gfa1-C562, but not other detected cysteine residues in either protein, undergo a significant decrease in oxidation in the absence of *SOD*1 (**Table S1J**). We also observed that in one case, Nbp35 (required for maturation of extramitochondrial Fe-S proteins), the site of dynamic oxidation is a functional metal binding site in the protein (**Table S1D**). Finally, other proteins not revealed by previous analyses were also evident, most notably Uba1, which is the sole enzyme necessary for activating ubiquitin in yeast. Uba1-C447, which is immediately adjacent to the ATP binding site for the protein, undergoes a large drop in oxidation in the absence of SOD1 – going from a maximum ∼89% in WT cells down to ∼28% in *sod1*Δ cells (**Figure S6B** and **Table S1J**). In comparison, C600, which is the site of ubiquitin attachment, undergoes very little change in oxidation upon deletion of *SOD1*. Collectively, these data provide an unparalleled look into the dynamic oxidation of specific cysteine thiols as well as protein abundance changes in response to the loss of *SOD1*.

## Discussion

Life in air necessitates that cells sense and detoxify ROS generated from aerobic metabolism. As a consequence, a number of enzymatic antioxidant defenses evolved to combat ROS, including O_2_^⸱-^-scavenging SODs and H_2_O_2_-scavenging thiol peroxidases and catalases. SODs are unusual “antioxidants” in that they catalyze the production of one ROS, H_2_O_2_, as a byproduct of detoxifying another ROS, O_2_^⸱-^. Rather curiously, prior studies found that increased expression of Sod1 reduces cellular H_2_O_2_ levels(2), hinting at additional unknown mechanisms underlying Sod1 antioxidant activity. Herein, we sought to understand if there were physiological roles for Sod1-derivied H_2_O_2_ in redox regulating antioxidant defenses and the thiol proteome. Indeed, we identified a novel redox circuit in which Sod1 senses the availability of O_2_ via metabolically produced O_2_^⸱-^ radicals and catalyzes the production of H_2_O_2_ that oxidatively inactivates GAPDH. GAPDH inactivation in turn promotes flux through the oxPPP to generate the NADPH required for aerobic metabolism and antioxidant defenses (30, 31). Moreover, using mass spectrometry-based redox proteomics approaches, we identified a larger network of proteins whose redox state, like GAPDH, is sensitive to Sod1 levels. Altogether, our results highlight a new mechanism for the antioxidant activity of Sod1 – namely that Sod1-derived H_2_O_2_ can stimulate NADPH production – and place Sod1 as a master regulator of the cellular redox landscape through two potential mechanisms; either by providing a direct source of thiol oxidizing H_2_O_2_ or by altering the NADPH/NADP+ redox balance through the Sod1/GAPDH signaling axis (**Figure 7**).

**Figure 7.**
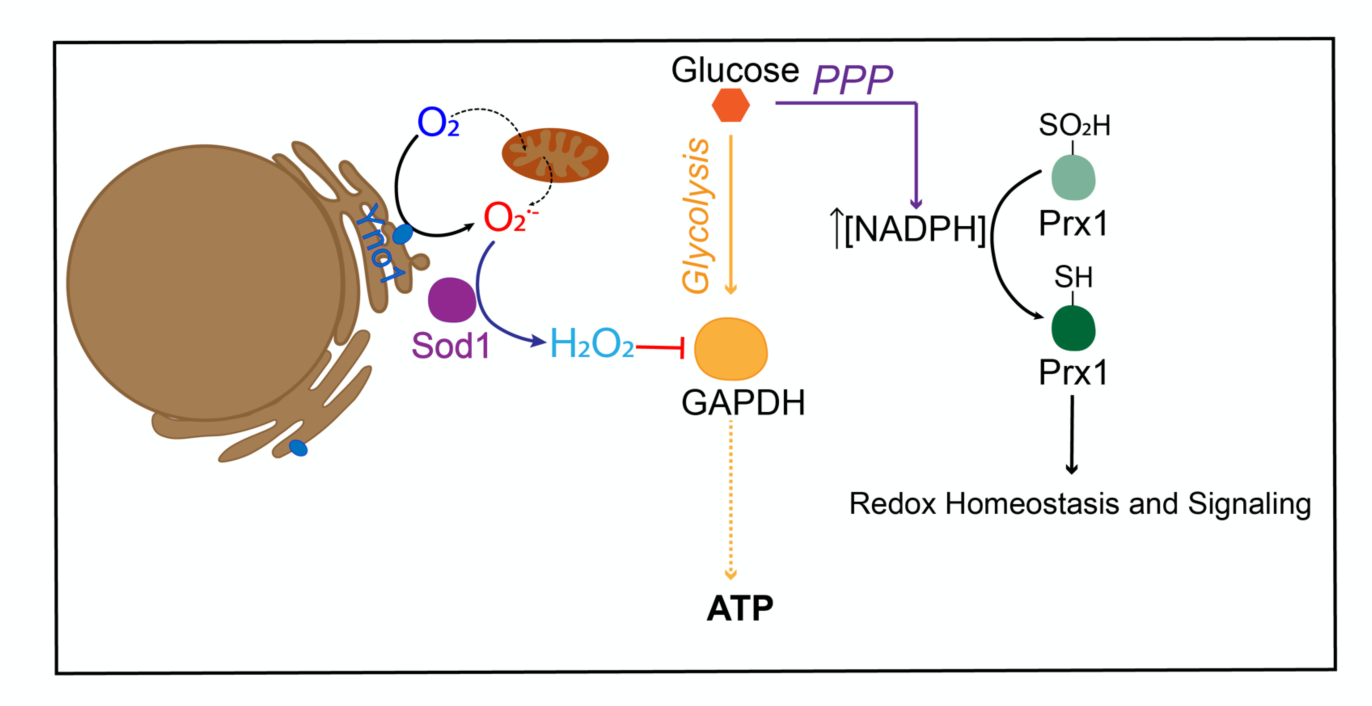
Proposed model for how Sod1 integrates O_2_ availability to regulate GAPDH oxidation and balance flux between glycolysis and the PPP to produce NADPH.

GAPDH catalyzes a rate determining step of glycolysis and is amongst the most abundant peroxide reactive enzymes in eukaryotes (23). Prior to our work, the oxidative inactivation of GAPDH and subsequent increase in NADPH via the PPP to combat redox stress was exclusively associated with high concentrations of exogenous oxidants (24, 30-33, 55). Our studies found that endogenous H_2_O_2_, specifically derived from Sod1, is responsible for the aerobic oxidation of GAPDH (**Figure 3D**). These results indicate that only a highly localized and privileged pool of peroxide is capable of physiological GAPDH oxidation and control of glycolytic flux. As GAPDH and Sod1 are amongst the most abundant soluble proteins in cells, we propose that there may be transient interactions between there proteins that facilitate Sod1-dependent GAPDH oxidation. While we could not detect the co-immunoprecipitation (Co-IP) of endogenous Sod1 and Tdh3 from yeast cells, we did observe that purified recombinant bovine Sod1 and yeast Tdh3 do interact as assessed by co-IP (data not shown). By extension, we suggest that signaling H_2_O_2_ is channeled between Sod1 and down-stream targets via protein-protein interactions, rather than through diffusion. This model for Sod1-based redox signaling complements other paradigms of peroxide signaling such as the transfer of oxidizing equivalents through thiol-disulfide exchange (56) or the “flood-gate” effect, which posits that a burst of H_2_O_2_ inactivates antioxidants like peroxiredoxins so that sufficient peroxide may diffuse far enough to transmit redox signals (57).

The aerobic oxidation of GAPDH is rate limited by Sod1 (**Figure 3E** and **3F**), suggesting that the oxidative inactivation of GAPDH and glycolytic flux may be dynamically regulated by Sod1 expression, maturation, which itself is O_2_ dependent, its interactions with GAPDH, or post translational modifications (PTMs) that regulate Sod1 localization and/or activity (58-61). With respect to Sod1 PTMs, it is tempting to speculate that the O_2_ sensing redox circuit we identified herein may be integrally linked to the nutrient sensing TOR pathway. TOR-dependent phosphorylation at Ser38 in yeast (Thr39 in humans) suppresses Sod1 activity in response to nutrient abundance to promote cell growth, whereas starvation promotes Sod1 activity (61). TOR/Sod1 signaling may be coupled to the redox regulation of GAPDH and glycolytic flux, thereby providing a means to integrate oxygen and nutrient availability to control metabolism. With respect to Sod1 expression, since many cancer cells over-express Sod1, it is conceivable that the Sod1/GAPDH signaling axis promotes cancer cell survival and drug resistance due to its potentiation of NAPDH production via the PPP (62). Thus, our work also sheds new light on the benefits of anti-Sod1 therapeutic interventions (61, 63-66). Indeed, we find that the breast cancer MCF-7 cell line exhibits greater levels of GAPDH oxidation relative to HEK293 cells, correlating with its higher levels of Sod1 expression.

The integration of O_2_ availability through the Sod1/GAPDH redox signaling axis is dependent on key sources of O_2_^⸱-^, including the yeast NADPH oxidase Yno1 and mitochondrial respiration (**Figure 2**). Given that Sod1 and GAPDH are primarily cytosolic enzymes, it is not surprising that we found Yno1, which is localized at the ER membrane and produces O_2_^⸱-^ on the cytosolic side, is required for transducing O_2_ availability to the O_2_^⸱-^ signal required for Sod1/GAPDH signaling (49). Indeed, Yno1 was previously found to regulate Sod1-mediated redox regulation of yeast casein kinase Yck1/2 signaling, which also occurs in the cytosol in proximity to the plasma membrane. However, it was rather surprising to find that mitochondrial respiration, as assessed in a respiratory incompetent yeast mutant lacking mitochondrial DNA, could also contribute O_2_^⸱-^ for Sod1 mediated GAPDH oxidation in the cytosol. Complex I (missing in *S. cerevisiae*) and III are the primary sources of electron leakage from the electron transport chain and produce O_2_^⸱-^ in the mitochondrial IMS (67). Yet, IMS targeted Sod1 does not mediate GAPDH oxidation (**Figure 1M** and **1N**). Thus, IMS O_2_^⸱-^, which is charged and membrane impermeable, must exit the mitochondria for Sod1/GAPDH signaling via voltage-dependent anion channels (VDACs) (68) or as membrane-permeable neutral hydroperoxyl radicals (HO_2_^⸱-^). Our results highlight how mitochondrial respiration may be an important sensor for O_2_, providing a source of O_2_^⸱-^ that can act as a retrograde signal to control extra-mitochondrial metabolism for adaptation to increasing *p*O_2_. The current study complements prior work that found mitochondrial respiration is a key source of ROS required for adaption to hypoxia via HIF signaling (69).

The 10^4^-rate enhancement of Sod1-catalyzed O_2_^⸱-^ disproportionation relative to the uncatalyzed reaction suggests that Sod1 acts as a redox amplifier that provides a localized burst of H_2_O_2_ within the vicinity of Sod1 for specific redox signaling events. Indeed, the O_2_-dependent redox regulation of GAPDH and casein kinase signaling are exclusively dependent on Sod1. These findings prompted us to determine if other redox targets of Sod1 exist. Indeed, mass spectrometry-based redox proteomics identified a number of proteins that, like GAPDH, are more oxidized in the presence of Sod1. Interestingly, enzymes associated with pathways orthogonal to glycolysis and the PPP, namely involved in amino acid biosynthesis, were prominently enriched over other groups (**Figures 6H, S5D** and **S5E**), and suggest that Sod1-dependent growth control may also be modulated through additional targets within these pathways that may have evolved Sod1-sensitive cysteine oxidation sites. On the other hand, all but one of these proteins (Nbp35) are oxidized at Cys residues not associated with a direct role in catalysis or function, raising the possibility that oxidation of such sites may function allosterically. For instance, the E1 ubiquitin activating enzyme Uba1 is differentially oxidized at C447 (**Table S1J, Figure S6B**), which is immediately adjacent to the ATP binding site, and the cysteine desulfurase Nfs1 is differentially oxidized at C199, which is adjacent to the site of pyridoxal phosphate cofactor attachment. Future work will probe the functional consequences of Sod1-dependent protein oxidation. We also identified proteins that are more oxidized in *sod1*Δ cells, which we interpret as arising due to oxidative stress associated with the loss of GAPDH/Sod1 signaling and the concomitant decrease in NADPH levels and/or O_2_^⸱-^ toxicity. It is worth noting that although past redox proteomics studies revealed much about potential targets of redox regulation, they typically involved treatment with exogenous oxidants mimicking pathological stress levels outside the physiological range (24, 70-73). Our study is notable in that we have identified redox targets of a physiological endogenous source of H_2_O_2_.

To what degree does Sod1 regulate thiol oxidation in animal models? A recent elegant redox proteomics study from Couchani and co-workers, termed “Oximouse”, describes tissue specific protein thiol oxidation in a mouse model (74). Since active Sod1 requires the formation of an intramolecular disulfide bond between Cys 147 and Cys 56, we plotted % Sod1-Cys147 oxidation as a proxy for mature active Sod1 against overall average Cys oxidation in each tissue and found a statistically significant linear correlation (r^2^ = 53; p = .04) (**Figure S7A**). To determine if this correlation between active Sod1 and overall thiol oxidation was unique to Sod1, we also plotted tissue Cys oxidation against the oxidation of active site Cys residues in other peroxide metabolizing enzymes, including peroxiredoxins 1 and 5 (PRDX1, PRDX5) or glutaredoxin 3 (GLRX3). Unlike Sod1, the activity of these other enzymes, as assessed by their active site oxidation, do not significantly correlate with overall Cys oxidation (**Figure S7B-S7F**). Moreover, we also found that the oxidation of other enzymes known to be redox regulated by Sod1, including tyrosine phosphatases PTP1N (PTP1B in humans) and PTPN11 (75, 76) and a GTPase involved in vesicular protein transport from the ER to the Golgi, Rab1*α* (77) (**Figures S7G-S7I**), exhibit a linear correlation between active Sod1 and their active site oxidation. Notably, there is no correlation with PRDX1 Cys oxidation, which is typically associated with the transmission of oxidizing equivalents to regulate redox signaling (56) (**Figures S7J-S7L**). Together, the data we extracted from the Oximouse study supports our principle findings from yeast that Sod1-derived peroxide regulates proteome-wide thiol oxidation.

Our studies have also finally explained a mystery regarding the role of Sod1 in antioxidant defense. Prior to the current work, the primary role of Sod1 in antioxidant defense was thought to be O_2_^⸱-^ scavenging. However, we previously found that the vast majority of Sod1, > 99%, was dispensable for protection against cell wide markers of O_2_^⸱-^ toxicity (15). We surmised that exceedingly low amounts of Sod1 were sufficient to protect against O_2_^⸱-^ because the targets of O_2_^⸱-^ toxicity are limited in scope, primarily Fe-S cofactor containing proteins. Indeed, all O_2_^⸱-^-related toxicity phenotypes arise from diminished activity of Fe-S enzymes and iron toxicity due to iron release from damaged clusters. Our work indicates that a broader role for Sod1 in oxidant defense is to promote the production of NADPH via the GAPDH/Sod1 signaling axis. As a major cellular reductant for numerous antioxidant systems, NADPH offers more expansive protection against redox stress than just defending against O_2_^⸱-^. Since GAPDH and Sod1 are amongst the most ancient and highly conserved enzymes in aerobic life, we propose that this new reported function of Sod1 to produce NADPH via GAPDH oxidation was a key requirement for O_2_ sensing and integration for adaptation to life in air.

## Acknowledgements

We wish to acknowledge Ms. Kathrin Ulrich for assistance with mPEG-mal labeling protocols, Prof. Valeria Culotta for the Sod1 antibody, Prof. Tobias Dick for the *tdh1Δ tdh2Δ tdh3*Δ strains, Prof. Dennis Thiele for the mitochondrial targeted Sod1 allele, and Prof. Breitenbach for the Yno1 plasmids. We acknowledge support from the U.S. National Institutes of Health (GM118744 to A.R.R. and M.P.T.) and the Blanchard Professorship (to A.R.R.).

## Author Contributions

C.M.A. co-designed and conducted the molecular genetics and biochemical experiments in yeast and human cell lines and analyzed the data. H.K. and A.P.J co-designed and performed mass spectrometry experiments and analyzed data. M.P.T. co-designed and supervised the mass spectrometry experiments and analyzed the data. A.R.R. co-designed and supervised experiments and analyzed data. C.M.A., M.P.T. and A.R.R co-wrote the paper, with input from all authors.

## Methods

### Chemicals, media components and immunological reagents

Dihydroethidium (Cat. # 50-850-563) was purchased from Thermo Fisher Scientific. Methoxypolyethylene glycol maleimide, 5kDa (Cat. # 63187) was purchased from Sigma Aldrich. SC and YPD dropout mixtures were purchased from Sunrise Science Products and VWR, respectively. The yeast GAPDH activity assay kit (Cat. # K680-100) and the NADP/NADPH quantification kit (Cat. #K347-100) were purchased from Bio Vision. Pierce anti-HA beads for immunoprecipitation of HA-tagged GAPDH was purchased from Thermo Fisher (Cat. # 88836) Rabbit polyclonal antibody (Cat. # G4595) and mouse monoclonal antibody against GAPDH (Cat. # G8795) were purchased from Sigma Aldrich. Rabbit polyclonal antibody against Peroxiredoxin-SO_3_ (Cat. # ab16830) was purchased from Abcam. Rabbit monoclonal antibody against HA tag (Cat. # 3724S) was purchased from Cell Signaling. Goat Anti-Rabbit (Cat. #89138-520) and anti-mouse CF680 (Cat. # 20067) secondary antibody were purchased from Biotium. Goat anti-rabbit DyLight 800 secondary antibody was purchased from Thermo Fischer Scientific (Cat. # SA5-35571). A previously described rabbit polyclonal antibody against Sod1 was obtained from the laboratory of Valeria Culotta (Johns Hopkins University)(16). Revert^TM^ 700 Total protein stain kit for protein normalization was purchased from (LI-COR).

### Yeast strains, plasmids, and growth

*S. cerevisiae* strains used in this study were derived from BY4741 (MATa, his3Δ1, leu2Δ0, met15Δ0, ura3Δ0). sod1::*KANMX4* strains were generated by knocking out *SOD1* using the previously described deletion plasmid pJAB002 (52). Expression constructs for wild type *SOD1* (pRS415-*SOD1*) and IMS localized SCO2-*SOD1* (pRS415-SCO2-*SOD1*), which are both driven by the native SOD1 promoter, were previously described and were provided by the laboratory of Professor Dennis Thiele (Duke University) (48).

The previously described *GAL1* driven *SOD1* expression plasmid (pAR1026) (15) was constructed by PCR amplification of the *SOD1* open reading frame from BY4741 genomic DNA with primers that introduced flanking 5′ and 3′ *SpeI* and *BamHI* sites, respectively. The *SOD1* amplicon was sub-cloned into the *SpeI* and sites of pRS316-*GAL1* (78) to generate pAR1026.

The ADH driven SOD1 expression construct (pCMA002) was constructed by PCR amplification of the *SOD1* open reading frame from *BY4741* genomic DNA using plasmids that introduced 5’ and 3’ flanking regions with the *BamHI* and *XhoI* restriction sites, respectively. The *SOD1* amplicon was cloned into the *BamHI* and *XhoI* cloning sites of p415ADH to generate pCMA002. The plasmid expressing *CCS1*-independent SOD1 mutant (pCMA010, p415ADH-*SOD1*-P144S) was generated using site directed mutagenesis on the pCMA002 plasmid (**Table S2**).

Expression constructs for Yno1 (p416YES2-*YNO1*) or EV (p416YES2-EV), which are driven by a strong galactose inducible promoter were previously described and were provided by the laboratory of Michael Breitenbach (University of Salzburg) (49). These constructs were cloned in the previously described sod1::kanMX4 strain expressing wild type *SOD1* (pRS415-*SOD1*).

A previously described 3Δ (312) TDH3 strain (BY4742 Δtdh3::*kanMX4* Δtdh1::*natNT2* Δtdh2::*hphNT1* + pDP8) expressing wild type *TDH3* (pDP8, p415TEF-TDH3) was provided by the laboratory of Tobias P. Dick (dkfz) (23). The resolving cysteine TDH3 mutant (p415TEF-*TDH3*-C154S) was generated via site directed mutagenesis (**Table S2**).

Yeast transformations were performed by the lithium acetate procedure (79). Strains were maintained at 30 °C on either enriched yeast extract-peptone based medium supplemented with 2% glucose (YPD), or synthetic complete medium (SC) supplemented with 2% glucose and the appropriate drop-out mixture to maintain selection. For all experiments, cells were streaked from -80 °C glycerol stocks onto solid agar media plates and pre-cultured in an anaerobic chamber (Coy laboratories) maintained with an atmosphere of 95% N2 and 5% H2. Anaerobically grown cells required supplementing SC media with 15mg/L of ergosterol and 0.5% Tween-80 (SCE) (80).

For typical experiments involving the IMS-targeted SCO2-*SOD1* expression plasmid, cells were cultured aerobically in SC-LEU, with 2% glucose. In all cases, cells were seeded at an OD_600nm_ ∼ .01 and cultured for 14-17 hours to a density of OD_600nm_ ∼ 1.0 at 30 °C in a shaking incubator (220 RPM). Following growth, cells were processed as described below for immunoblotting and enzyme assays. For experiments involving the *CCS1*-independent mutant *SOD1*-P144S, cells were cultured aerobically or anaerobically in previously degassed SCE-LEU with 2% glucose. Anaerobic cultures were seeded at an OD_600nm_∼ .08 and cultured for 14-17 hours to a density of OD_600nm_ ∼ 1.0 at 30 °C shaking in the anaerobic chamber (200 RPM). For this experiments, Sod1 activity and/or expression was assessed as described below. For experiments involving the titration of *YNO1* and *SOD1* using the GAL-driven *YNO1 and SOD1* expression plasmids, cells were cultured aerobically in SC-URA with 2% raffinose and the indicated galactose concentrations.

Human embryonic kidney HEK293 cells were obtained from the laboratory of Loren D. Williams (Georgia Institute of Technology). Cells were cultured in Dulbecco Modified Eagle’s Medium (DMEM, VWR), with 4.5g/L glucose and without L-Glutamine or Sodium Pyruvate, supplemented with 10% fetal bovine serum (FBS). *SOD1* silencing was accomplished using 50µM SOD1 silencer (Ambion, Cat. # 4390824) or scrambled control (Ambion, #Cat AM4611). Cells were transfected in Opti-MEM (Fisher) with Lipofectamine 2000 (Invitrogen) for 72h. MCF7 cells were cultured in DMEM without Phenol Red or L-glutamine (VWR).

All experiments were conducted using biological replicates arising from duplicate, triplicate or more independent cultures of multiple clones. All of the data has re-produced on multiple occasions in independent experimental trials.

### TCA precipitation and thiol alkylation with mPEG-mal in yeast cultures

10 mL aerobic or anaerobic overnight yeast cultures were quenched at a density of OD_600nm_ mL^-1^ ∼ 1 by adding cold 100% trichloroacetic acid (TCA) to a final concentration of 10% (v/v). The TCA-stopped cultures were incubated on ice for 1h before pelleting and washing in 20%TCA. The final pellet was stored at −80°C. For experiments involving H_2_O_2_ treatment, H_2_O_2_ was added to a final concentration of 1 mM to yeast cultures and incubated for 2 minutes while shaking prior to TCA precipitation.

Pellets were thawed on ice and lysed at 4°C in TCA lysis buffer (12.5% v/v TCA in 1 mM Tris-HCl, pH = 8.0, 25mM NH_4_Ac,1mM Na_2_EDTA pH 8.0) using half pellet volume of zirconium oxide beads and a beat beater on a setting of 8 for 3 minutes, twice (Bullet Blender, Next Advance) (46). The TCA lysate was transferred to a fresh tube by poking a hole 45° from the cap using a hot needle and pelleted, washed in cold acetone (−20°C) and dried. The dried pellet was resuspended in 200 µL degassed resuspension buffer (6M urea, 10mM EDTA, 20mM Tris, 0.5% w/v SDS, 10µM neocuproine, pH 8.5) containing 1mM PMSF and a protease inhibitor cocktail (GBiosciences) in the anaerobic chamber (Coy laboratories). Lysate protein concentrations were determined by the Pierce^TM^ BCA protein assay kit (Thermo Scientific).

Alkylation of free thiols was accomplished by diluting the lysate to a protein concentration of 0.18 µg/µL in resuspension buffer with 7.5 mM mPEG-mal (dissolved in DMSO) or DMSO and incubated for 1h shaking in the anaerobic chamber protected from exposure to light. The excess mPEG-mal was discarded using PD SpinTrap G-25 columns (GE Healthcare) and the resulting lysates were electrophoretically separated by SDS-PAGE in 14% tris-glycine gels (Invitrogen). Anti-GAPDH (1:4000) polyclonal rabbit antibody and a goat secondary antibody conjugated to a 680nm emitting fluorophore were used to visualize GAPDH in the immunoblot. All gels were imaged on a LiCOR Odyssey Infrared imager. Immunoblot quantitation was conducted using the LIC-COR Odyssey imager software (Image Studio Lite). %GAPDH oxidation was calculated dividing the intensity of the bottom band (unlabeled oxidized GAPDH) by both top and bottom bands (total detected GAPDH, labeled plus unlabeled).

### Enzyme assays

SOD activity analysis was performed by native PAGE and nitroblue tetrazolium staining as described previously (15, 16, 81, 82) on exponential phase aerobic and anaerobic cultures grown to a density of OD_600nm_ ∼ 1.0 in degassed SCE-LEU, 2% glucose media. Yeast cells were washed in ultrapure H_2_O, resuspended in cold (4°C) lysis buffer (10mM sodium phosphate, 50 mM sodium chloride, 5 mM EDTA, 0.1% Triton X-100, 1 mM PMSF and protease inhibitor cocktail). Lysis was achieved at 4°C using half pellet volume of zirconium oxide beads and a beat beater as described before (15). Lysate protein concentrations were quantified by the Bradford method (Bio-Rad) and 15 µg of each protein sample was loaded and separated in 14% native PAGE gels. Sod1 activity was visualized by staining the gel with SOD activity staining solution (2.43 mM nitro blue tetrazolium chloride, Sigma, 0.14M riboflavin-50-phosphate, Sigma, 45mM dipotassium phosphate buffer, 4.5mM monopotassium phosphate buffer) with 28mM TEMED (Bio-Rad) for 60 minutes at room temperature in darkness. After the incubation, gels were rinsed with water twice and exposed to light.

GAPDH activity was measured using the GAPDH activity assay kit (Bio vision) according to the manufacturer’s specifications. For this purpose, *sod1*Δ cells expressing an empty vector (pRS415) or SOD1 (pRS415-SOD1) were grown in 10mL SC-LEU 2% glucose cultures to a density of OD_600nm_=1.0. Cells pellets were washed in ultrapure water and lysed in cold degassed lysis buffer as described for the SOD activity analysis. Increasing concentrations of protein lysate within a linear detection range were diluted in assay buffer to a final volume of 50 µL and mixed with 50 µL of reaction mix. Activities were monitored by spectrophotometrically measuring the generation of 1,3-bisphosphoglycerate (BPG) at 450 nm over the course of 60 minutes using a Biotek Synergy Mx multi-modal plate reader.

ATPase activity was assayed in membrane fractions with or without a 5 min preincubation with 50 mM vanadate. ATPase activity was determined by quantitating the phosphate released during vanadate-sensitive ATP hydrolysis by molybdate reactivity (83), and activity was normalized to WT cells.

### NADPH measurements

NADP/NADPH ratio measurements were conducted using the NADPH kit (Bio vision) according to the manufacturer’s specifications. Briefly, yeast 10 mL cultures were split in half after reaching an OD_600_/mL∼1.0, and 5 OD_600_ were TCA precipitated, as previously described, to assess peroxiredoxin oxidation (Prdx-SO_3_) and the other 5 OD_600_ were harvested and lysed in the NADP/NADPH extraction buffer provided by the manufacturer. Half the lysate was incubated on ice to assess total [NADP] and [NADPH], and the other half was heated for 30 minutes at 60°C in a heat block to assess [NADPH]. Increasing concentrations of protein lysate within a linear detection range were diluted in assay buffer to a final volume of 50µL and mixed with 100µL of reaction mix and 10µL of Developer. NADPH was monitored after 1h incubation every 20 minutes for 4h at 450nm using Biotek Synergy Mx multi-modal plate reader.

### Superoxide measurements

Superoxide levels were measured by monitoring the fluorescence of DHE stained cells (λex =485 nm, λem =620 nm) as previously described (15). Briefly, 1 x10^7^ cells were harvested from triplicate cultures, resuspended and incubated in 500µL filter-sterilized 1xPBS solution containing 50µM DHE for 20 minutes at 30°C in the dark, washed twice in 1xPBS. The fluorescence was recorded in a Biotek Synergy Mx multi-modal plate reader.

### Growth test

Growth tests of WT and *tdh3*Δ aerobic and anaerobic cultures were performed growing three biological replicates anaerobically overnight and diluting each biological replicate to 0.15 OD_600_/mL in duplicate 10mL cultures the next morning. Each replicate was cultured aerobically or anaerobically for 12h. OD_600_/mL readings were taken every hour or 2h from aerobic and anaerobic cultures, respectively, using a UV/vis spectrophotometer (Agilent). For peroxide sensitivity tests, WT and *tdh3*Δ aerobic cultures were grown overnight and diluted to 0.15 OD_600_/mL with increasing concentrations of H_2_O_2_ (0, 100µM, 250µM, 500µM, 1mM). Readings were taken every hour using a UV/vis spectrophotometer (Cary 60, Agilent).

### Immunoprecipitation

Immunoprecipitation of HA tagged GAPDH (TDH3-HA) was performed using Pierce anti-HA beads (Thermo Fisher). Briefly, 100mL cultures of 3Δ (312) TDH3 strain expressing TDH3-HA or WT-TDH3 were grown overnight to an OD_600_/mL ∼ 1.0 in 500mL Erlenmeyer flasks. The corresponding lysates, at a final concentration of 3 mg/mL, were incubated with 50µL of anti HA beads for 2h rotating end over end at room temperature (RT), after the beads had been washed with TBS-T (1x TBS containing 0.05% Tween-20). Next, after the remaining lysates were discarded, and after several washes with TBS-T, the beads with the immunoprecipitated TDH3-HA (IPed) were incubated with increasing concentrations of bovine recombinant Sod1 in lysis buffer, corresponding to 0.01, 0.1 and 1 equivalent of Sod1 for 30 minutes at RT. After several washes with TBS-T, the IPed material was eluted by incubating the beads with 50µL of 0.1M Glycine buffer, pH 2.0, for 10 minutes and the resulting eluate was neutralized with 7.5µL of 1M Tris-HCl buffer, pH 8.5.

### HEK293 and MCF7 TCA precipitation and thiol alkylation with mPEG-mal

For assessing Sod1-dependent GAPDH oxidation in HEK293 cells, cells were silenced by adding 50µM SOD1 silencer (Ambion, Cat. # 4390824) or scrambled control and Lipofectamine 2000 (Invitrogen) in Optimem to cells seeded in 6 well plates and incubated for 72h. GAPDH oxidation was assessed by lysing cells via TCA precipitation 72h-post transfection where 100% cold TCA was added to the collected cell suspension to a final concentration of 10% and incubated for 1h on ice. After TCA lysis, cells were pelleted and washed in cold acetone and dried. Dried pellet resuspension, thiol alkylation and immunoblotting were carried out as described in the section on TCA precipitation and thiol alkylation in yeast cultures, using a concentration of 0.5µg/µL lysate protein for thiol alkylation.

SOD1 silencing was assessed via immunoblotting from the cell lysates used for mPEG-mal labeling using a Sod1 a polyclonal rabbit antibody (1:1000) and a goat secondary antibody conjugated to a 680nm emitting fluorophore. GAPDH was used as the loading control and was visualized using a GAPDH mouse monoclonal antibody (1:4000) and a goat antibody conjugated to an 800nm emitting fluorophore. Silencing efficiency was calculated by normalizing Sod1 expression (measured normalizing the Sod1 band intensity to the GAPDH band intensity) from siSOD1 or siCTRL treated cells to the average Sod1 expression value from siCTRL treated cells.

For assessing GAPDH oxidation via exogenous peroxide addition, ∼10^5^ cells were seeded in a 6-well plate in a reduced-serum media (Optimem, Gibco), grown for 24h and treated with 100µM H_2_O_2_ dissolved in Optimem (37°C) for 2 minutes prior to cell harvesting and TCA lysis. As a control, cells were incubated with 2mL Optimem containing the same volume in H_2_O.

For assessing GAPDH oxidation in MCF-7 cells, cells were seeded in a 6 well plate and harvested and lysed via TCA precipitation 24h after the cells were seeded, as previously described.

### Quantification and Statistical analysis

Data and statistical analysis were performed using GraphPad Prism, Pymol and Studio Lite software. P values were calculated using two-tailed Student’s t-test for pairwise comparisons, One-way Anova and Dunett’s post-Hoc test for multiple comparisons or Two-way Anova for multiple comparisons with more than one variable. The P value for linear regression analysis was calculated using GraphPad Prism.

### SILAC labeling and cell lysis

Three biological replicates were analyzed each for WT and *sod1*Δ cells. For each replicate, *lys1*Δ and *lys1Δ SOD1::KANMX4* single colonies were inoculated in 3mL starter cultures and grown to saturation in synthetic complete (SC). The following day, starter cultures were diluted to ∼0.1 OD_600_ in 10mL dropout SC media to reach 0.8 OD_600_/mL and then seeded at an OD_600_ 0.0075 in 50mL SC media containing heavy (WT) or light (*sod1*Δ) lysine, respectively. Culture growth was stopped after ∼10 doublings by adding 100% cold TCA to a final concentration of 10%. The cells were incubated on ice in 10%TCA for 1h and pelleted for 10 minutes at 4,000 rpm, 4°C. The cell pellets were then washed in 20% TCA and pelleted for 10 minutes at 16,000 G, 4°C. After discarding the supernatant, the pellets were stored at −80°C.

To prepare extracts, the SILAC-labelled cell pellets were separately thawed on ice and resuspended in 20% TCA followed by vortexing with glass beads for 10 min at 4 °C. Lysates were transferred to new centrifuge tubes followed by centrifugation at 16,000 × g for 10 min at 4 °C. Pellets were washed with ice cold acetone followed by centrifugation at 16,000 × g for 5 min at 4 °C. Pellets were resuspended in freshly prepared 6 M urea, 200 mM Tris pH 8.0, 2% SDS, 10 mM EDTA, 10 µM neocuproine. Protein concentrations for both WT and sod1Δ resuspensions were determined by DC protein assay (Bio-Rad Laboratories) and mixed equivalently generating 400 µg total.

### iodoTMT labeling

The combined SILAC-labeled protein samples (heavy/light) were labeled using iodoTMT sixplex Isobaric Label Reagent Set (Thermo Scientific) according to the manufacturer specifications. Each bio-replicate was labeled with two different iodoTMT labels (either 126/129, 127/130, or 128/131). For pre-reduction iodoTMT-labeling, each replicate was labeled overnight followed by removal of unreacted TMT label by chloroform-methanol precipitation as previously described (84). Following the first labeling reaction, oxidized thiols were reduced in 50 mM DTT at 37 °C for 1 hr followed by excess DTT removal by chloroform-methanol precipitation. Post-reduction iodoTMT-labeling was carried out as before with overnight incubation followed by the unreacted TMT label reagent removal. Each replicate was washed with ice cold acetone followed by resuspension in 100 mM Tris pH 8.0, 1 M ABC, 6 M urea and incubation at 37 °C for 1 hr. Lys-C protease (Thermo) was added to each replicate (1:20; enzyme:protein) followed by overnight incubation at 37 °C with shaking at 600 rpm.

### High pH RPLC fractionation

SILAC-TMT labeled peptides were pre-fractionated by high pH reverse phase liquid chromatography using an Agilent 1100 HPLC system with Gemini C18 reverse phase column (2 × 150 mm, 5 µM, 110 Å; Phenomenex Inc.) with solvents A (10 mM ammonium formate) and B (10 mM ammonium formate in 90% acetonitrile) adjusted to pH 10 using ammonium hydroxide. A gradient method (from 0% to 70% B in 150 min, then from 70% to 0% in 20 min) was applied at a flow rate of 100 µl/min. Fractions were collected every 5 min and frozen at −80 °C followed by lyophilization by CentriVap.

### LCMS

Peptide fractions were analyzed by LCMS using a Q-Exactive Plus orbitrap mass spectrometer equipped with Dionex UltiMate 3000 LC system (Thermo). The lyophilized fractions were resuspended in 0.1% FA in 5% ACN and loaded onto a trap column, 75 µm I.D. x 2 cm, packed with Acclaim PepMap100 C18, 3 µm, 100Å (set of 2) nanoViper and resolved through an analytical column, 75 µm I.D. x 15 cm, packed with Acclaim PepMap RSLC C18, 2 µm, 100Å, nanoViper at a flow rate of 0.3 µl/min with a gradient solvent A (0.1% FA in 2% ACN) and a gradient solvent B (0.1% FA in 90% ACN) for 150 minutes. MS analysis was conducted in a data dependent manner with full scans in the range from 200 to 1800 m/z using an Orbitrap mass analyzer at a mass resolution of 70,000. The top fifteen most intense precursor ions were selected for MS2 with the isolation window of 4 m/z. Isolated precursors were fragmented by high energy collisional dissociation (HCD) with normalized collision energies (NCE) of 28, 31, and 33 for each analytical replicate. This led to the acquisition of 3 analytical replicates for each bio replicate for a grand total of 9 detection attempts per peptide fraction.

### MS data analysis

For SILAC data evaluation for protein expression, a total of 33,289 proteins of 3 biological replicates with 3 different NCEs of 28, 31, and 33 analyzed by PD 2.0 (**Table S1K**). Of these, 32,630 proteins were further evaluated excluding transposable elements. Protein SILAC ratios (*sod1*Δ_Light_/ WT_Heavy_) were calculated as the ratio of integrated MS1 peak areas for light and heavy peptides. Statistical significance in the difference between WT and *sod1*Δdata were determined using Response Screening (JMP 15.0) wherein up to 9 average responses per protein per yeast strain (18 total) were possible. Cases in which only one channel of a SILAC pair was present (heavy or light) were processed separately are reported as +/-infinite change in protein abundance.

For iodoTMT data evaluation a total of 307,894 peptides represented by 2,398,648 PSMs were identified using PD 2.0 (**Table S1L**). Of these, 35,476 unique peptides (259,936 PSMs) corresponded to iodoTMT-labeled cysteine-containing peptides, again with transposable elements excluded. These data were processed at the peptide level to calculate percent cysteine oxidation level by dividing the reporter ion signal of TMT2 by the sum of signals from TMT1 and TMT2, producing an oxidation percentage for WT_Heavy_ and *sod1*Δ_Light_ peptide isoforms. Statistical significance in the difference between WT and *sod1*Δ data were again determined using Response Screening (JMP 15.0) for oxidation percentages measured between the two strains wherein up to 9 average responses per protein per yeast strain (18 total) were possible. Cases in which only one SILAC peptide was present were processed separately. To examine overall differential oxidation levels of proteins, the cysteine coverage of a protein (# detected Cys / # total Cys in a protein) was calculated using the non-redundant UniProt Yeast (*BY4741*) proteome database. Custom scripts necessary to organize data in the context of cysteine native position and to determine relative cysteine detection coverage per protein were written in Perl 5.3.

### Bioinformatics

SILAC-iodoTMT proteomics data were further analyzed using PANTHER, an ontology-based database equipped with data analysis tools. GO enrichment of biological processes, molecular functions, and cellular components was performed for proteins with greater than +1 or less than −1 log_2_ SILAC ratio (sod1Δ_Light_ / WT_Heavy_), representing differential protein abundance. Separately, similar GO enrichment analysis was performed for proteins with observed TMT-labeled cysteine residues falling greater than 5% (or 10%) or less than −5% (or −10%) differential oxidation (*sod1*Δ – WT). Enriched terms with statistical significance above the 95% confidence level were considered further. GO enrichment bins were defined by first identifying GO terms with fold enrichment values that were statistical outliers (>90^th^ percentile), which resulted in well-defined bins reported in **Figures 6E**, **6H** and **Tables S1B, S1H** The mean fold enrichment for the GO Terms embedded with each bin was reported to allow comparison of fold enrichment values within each bin. Evaluation of cysteine oxidation site bio-process outliers (**Figure 6I**, **6J**, **S6A**, and **Table S1I**) was performed as follows. First, SGD GO analysis and EBI QuickGO were utilized to generate the list of bio-process GO terms for proteins undergoing statistically significant changes in oxidation (99 proteins in total). Second, GO terms were searched individually by AmiGO 2 to retrieve proteins in each of the GO terms filtered by *S. cerevisiae* S288C (see **Figure S6**). Third, the resulting list was filtered against proteins where cysteine oxidation in WT and sod1D cells could be quantified, and then further filtered to retrieve bio-processes in which any of the 99 significant ox changers were found to be a statistical outlier from other cysteine residues in the bio-process. Outlier responses of any cysteine oxidation change were highlighted by determining the distribution of observed responses within the bio-process. Outliers defined by this process were used to highlight specific proteins shown in **Figures 6I** and **6J**.

Pathway analysis of significant ox changers was further analyzed via the Kyoto Encyclopedia of Genes and Genomes (KEGG) Brite and Pathway analysis tools (see **Figure 6G**; **Tables S1F**, **S1G**). Identification of functional cysteine residues was achieved using UniProt curated data with restriction to “Binding Site”, “Active Site”, “DNA Binding”, or “Metal Binding”.

## Supplemental Information Title and Legends

**Figure S1.**
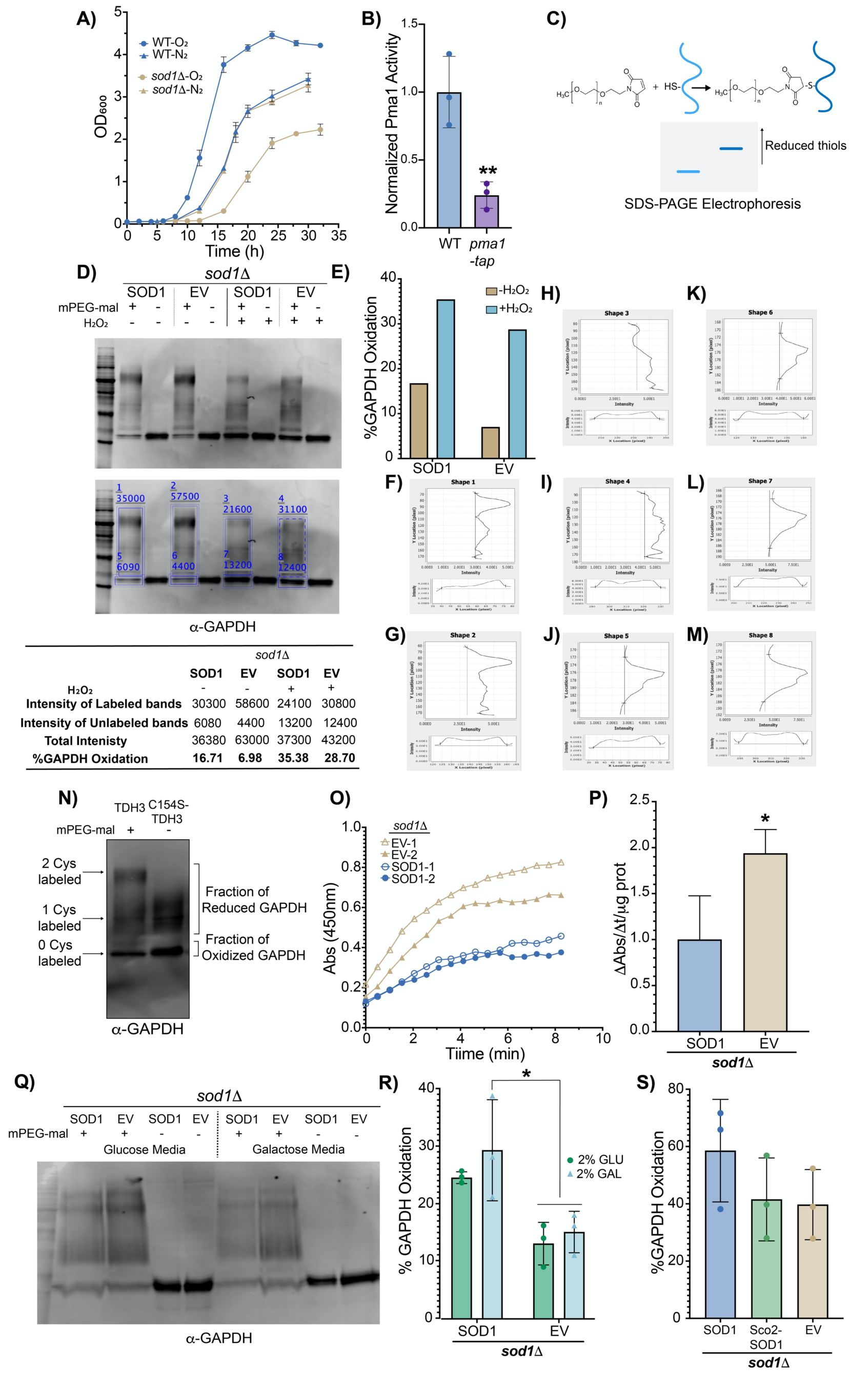
Development and validation of key strains, reagents, and techniques to establish Sod1-dependent GAPDH oxidation; related to Figure 1. **(A)** Aerobic (O_2_) and anerobic growth (N_2_) of WT and *sod1*Δ cells. Data represent the average ± SD from duplicate cultures. **(B)** Normalized Pma1 activity in WT cells and cells expressing a hypomorphic allele of *PMA1* (*pma1-tap*). Data represent the average ± SD from triplicate cultures and is normalized to the average activity of WT cells. **(C)** Schematic representation of Cys-mPEG-mal adducts upon covalent modification with methoxypolyethylene glycol maleimide (mPEG-mal) and the corresponding electrophoretic shift. **(D)** Representative immunoblot analysis of GAPDH-mPEG-mal adducts in *sod1*Δ cells expressing yeast Sod1 (SOD1) or empty vector (EV) cultured in 2% GLU and treated with 1mM H_2_O_2_ (+ H_2_O_2_) or H_2_O (-H_2_O_2_) for three minutes prior to treatment with mPEG-mal. Immunoblot quantitation was conducted using the LIC-COR Odyssey imager software (Image Studio Lite) (middle), yielding the values exhibited in the blue squares and the bottom table. The percentage of GAPDH oxidation is displayed in the bottom table and obtained dividing the intensity of unlabeled bands by the sum of the intensity of labeled and unlabeled bands. **(E)** % GAPDH oxidation, as assessed by quantifying the ratio of mPEG-mal labelled GAPDH to total GAPDH, in the indicated strains. **(F to M)** Peak profiles (Y location, top; X location, bottom) corresponding to the blue shapes displayed in panel D (middle) used to identify the intensity of the immunoblot bands accounting for background. The quantitation is performed using the Studio Lite software. **(N)** Representative immunoblot analysis of GAPDH-mPEG-mal adducts in *tdh1Δtdh2Δtdh3*Δ cells expressing yeast GAPDH (TDH3, left) or C154S-TDH3 (right). The arrows indicate the TDH3 or TDH3-C154S-mPEG-mal adducts. **(O-P)** Measurements of time-dependent (**O)** or average **(P)** GAPDH enzymatic activity in *sod1*Δ cells expressing SOD1 (blue) or EV (gold). Data represents the average ± SD from 2 **(O)** and 4 **(P)** independent trials. **(Q-R)** Assessment of GAPDH oxidation in *sod1*Δ cells expressing yeast Sod1 (SOD1) or empty vector (EV) cultured in 2% glucose (GLU) or 2% galactose (GAL). Representative immunoblot analysis of GAPDH-mPEG-mal adducts **(Q)** and absolute %GAPDH oxidation from multiple trials **(R)** in the indicated strains. Data represents the average ± SD from 3 **(S)** Assessment of GAPDH oxidation in *sod1*Δ cells expressing yeast Sod1 (SOD1), mitochondrial IMS targeted Sod1 (Sco2-SOD1), or empty vector (EV). The statistical significance is indicated by asterisks using two-tailed Student’s t-test pairwise comparison (panels B and P) or two-way ANOVA for multiple comparisons with Dunett’s post-hoc test for the indicated pairwise comparison (panel R). *p<0.05, **p<0.01, ***p>0.001, ****p<0.0001, n.s.= not significant.

**Figure S2.**
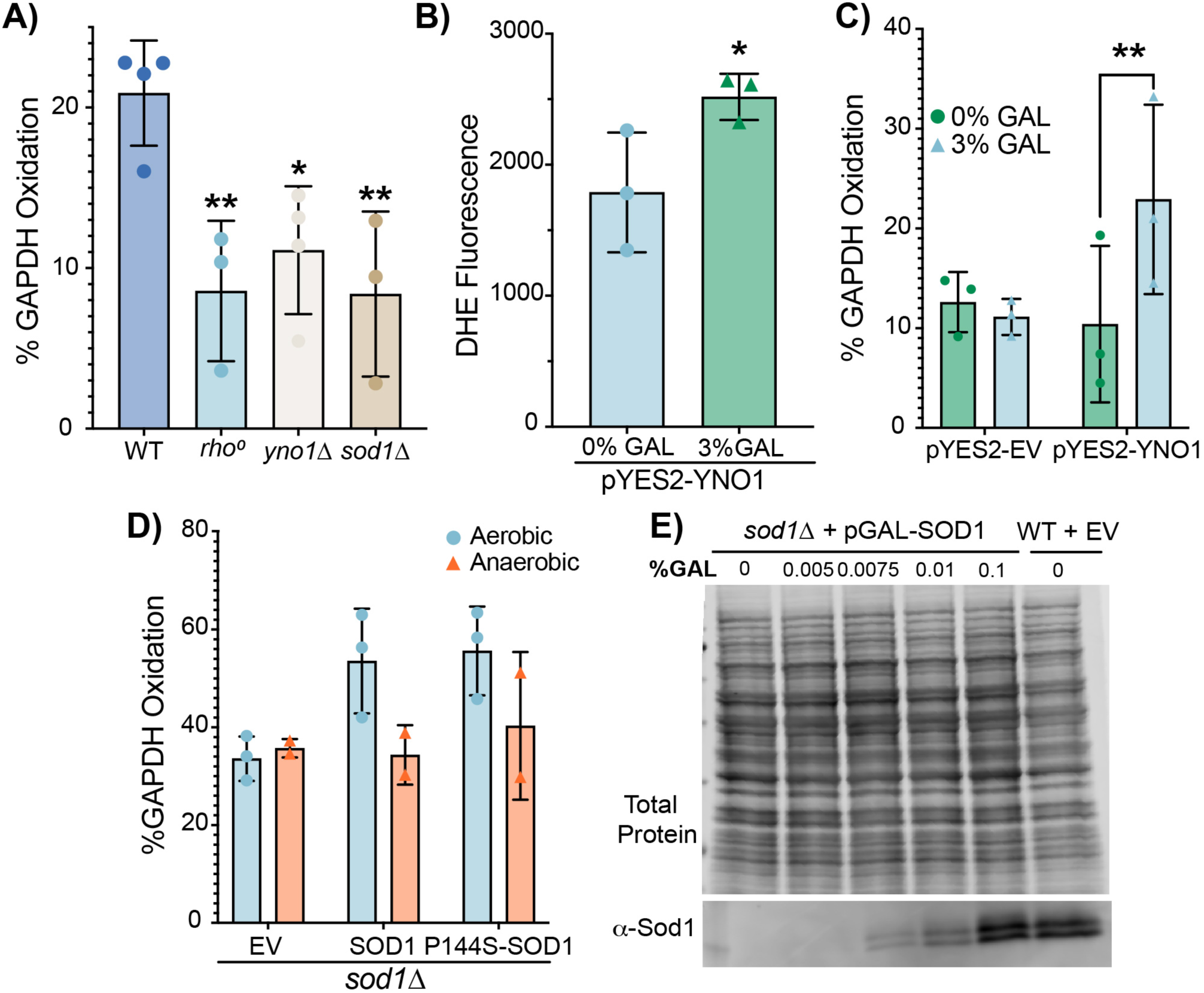
The effect of Yno1, respiration, O_2_, and Sod1 expression on % GAPDH oxidation; related to Figures 2 and 3. **(A)** % GAPDH oxidation assessed by mPEG-mal labeling of GAPDH in WT, *rho^0^*, *yno1*Δ and *sod1*Δ cells cultured in 2% GLU (Fig. 2A) from multiple trials. Data represents the average ± SD from 3 independent trials. **(B-C)** DHE-detectable superoxide **(B)** and percentage GAPDH oxidation **(C)** in in WT cells expressing GAL1-driven *YNO1* (pYES2-YNO1) or empty vector (pYES2-EV) cultured in 2% raffinose, supplemented with non-inducing (0%) or inducing (3%) GAL concentrations from multiple trials. Data represents the average ± SD from 3 independent cultures. **(D)** Assessment of the Sod1-dependence on the aerobic oxidation of GAPDH. Percentage GAPDH oxidation in aerobic or anaerobic *sod1*Δ cells expressing empty vector, WT SOD1, or the P144S SOD1 mutant (Fig. 3C). Data represents the average ± SD from two or three independent trials. **(E)** Representative immunoblot of Sod1 from *sod1*Δ cells expressing *GAL1* driven *SOD1* or WT cells transformed with empty vector (EV) cultured in the indicated concentrations of galactose (GAL). The statistical significance is indicated by asterisks using two-tailed Student’s t-test pairwise comparison (panel B), one-way ANOVA for multiple comparisons with Dunett’s post-hoc test for the indicated pairwise comparison (panel A) or two-way ANOVA for multiple comparisons with Dunett’s post-hoc test for the indicated pairwise comparison (panel C and D). *p<0.05, **p<0.01, ***p>0.001, ****p<0.0001, n.s.= not significant.

**Figure S3.**
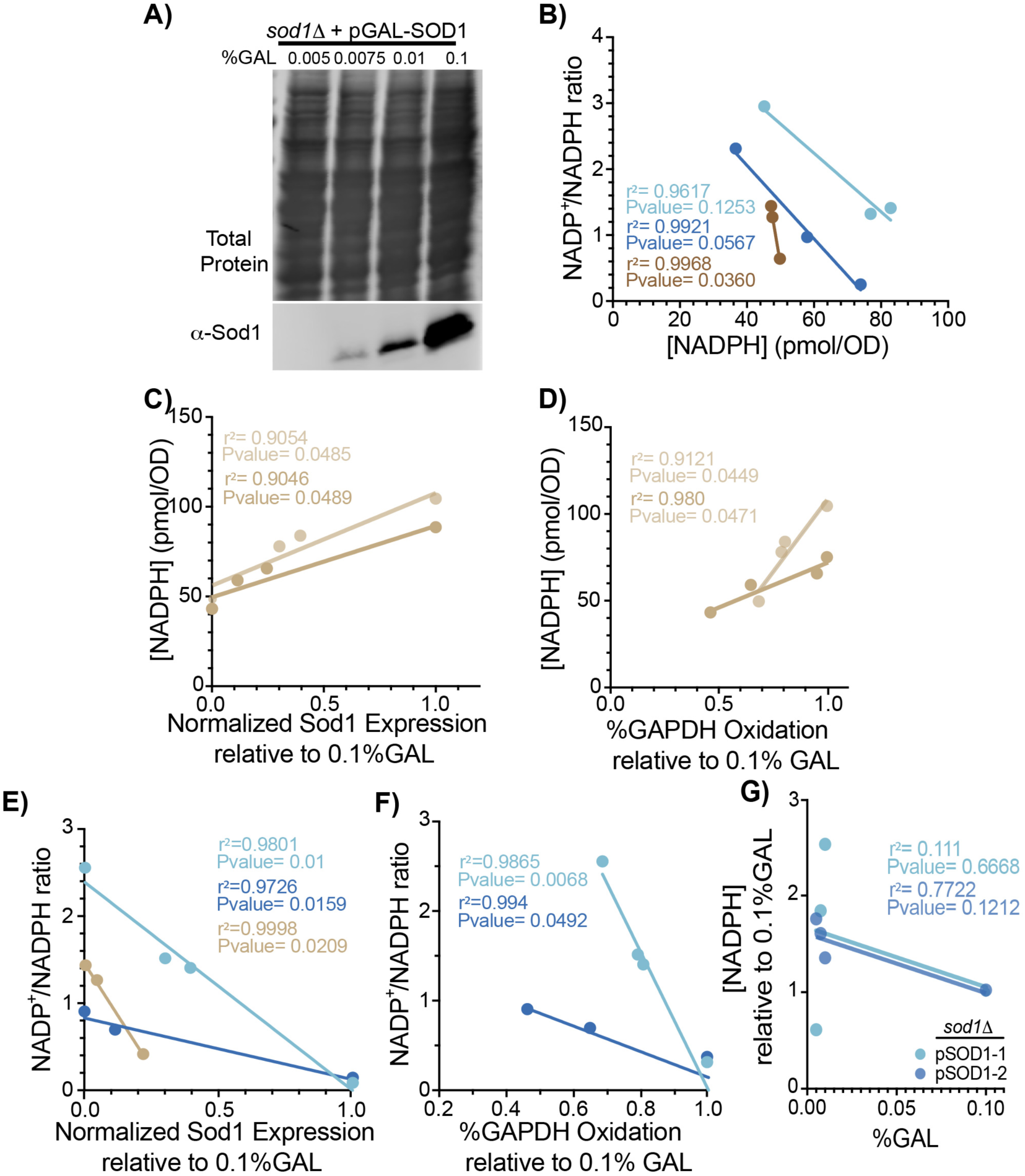
Titration of Sod1 results in a dose-dependent increase of [NADPH] and NADP^+^/NADPH ratio; related to Figure 4. **(A)** Immunoblot analysis of Sod1 expression and total protein quantification in *sod1*Δ cells expressing GAL-driven *SOD1* cultured with increasing concentrations of galactose (GAL) (0.005%, 0.0075%, 0.01% and 0.1% GAL). **(B)** The NADP+/NADPH ratio inversely correlates with cellular [NADPH]. The linear regression analysis in three independent trials gives coefficient of determinations (r^2^) of .96, .992 and .9968 and p-values of .12, .057 and .0036, respectively. **(C-D)** The [NADPH] correlates with normalized Sod1 expression **(C)** and normalized GAPDH oxidation **(D)**. Each color indicates two individual trials used for both correlation with Sod1 expression and GAPDH oxidation. The linear regression analysis in panel **C** gives coefficient of determinations (r^2^) of .9054 and .9046 and p-values of .0485 and .0489, respectively. The linear regression analysis in panel **D** gives coefficient of determinations (r^2^) of .9121 and .98 and p-values of .0449 and .0471, respectively. **(E-F)** The NADP+/NADPH ratio correlates with normalized Sod1 expression **(E)** and normalized GAPDH oxidation **(F)**. Each color indicates individual trials used for both correlation with Sod1 expression and GAPDH oxidation. The linear regression analysis in panel **E** gives coefficient of determinations (r^2^) of .9801, .9726 and .998 and p-values of .01, .0159 and .0209, respectively. The linear regression analysis in panel **F** gives coefficient of determinations (r^2^) of .9865 and .994 and p-values of .0068 and .0492, respectively. **(G)** Increasing concentrations of GAL do not positively correlate with [NADPH] in *sod1*Δ cells expressing a non-GAL regulated allele of yeast *SOD1* (p*ADH1*-SOD1). The linear regression analysis in panel **G** gives coefficient of determinations (r^2^) of .11 and .772 and p-values of .067 and .12, respectively.

**Figure S4.**
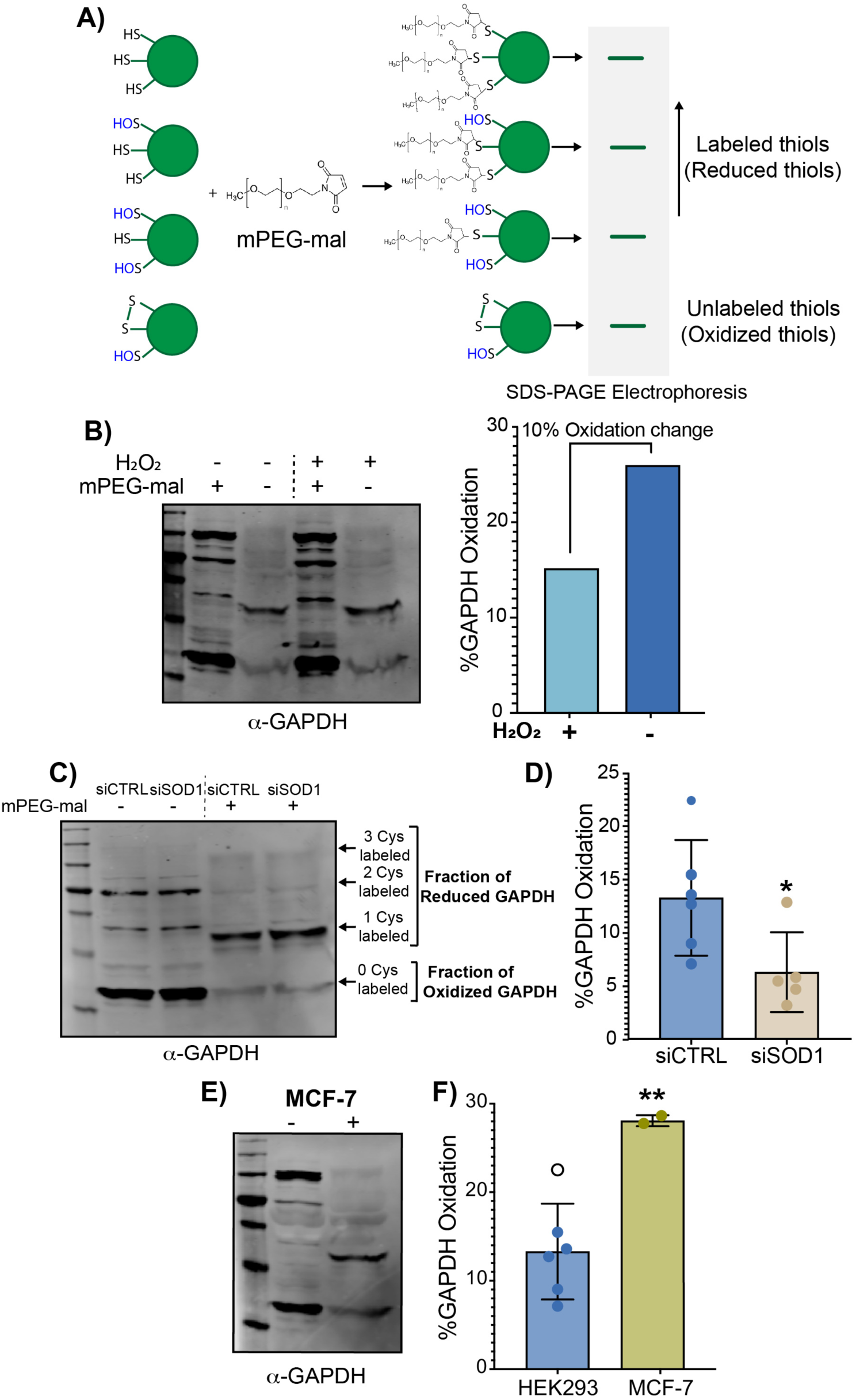
Validation of H_2_O_2_ and Sod1-mediated GAPDH oxidation in human cell lines; related to Figure 5. **(A)** Schematic representation of all possible redox-dependent GAPDH (green spheres) Cys-mPEG-mal adducts and their respective electrophoretic mobilities. **(B)** Representative immunoblot analysis of GAPDH oxidation as assessed by mPEG-mal labeling from human embryonic kidney HEK293 cells that have been treated with 100µM 100µM H_2_O_2_ (+H_2_O_2_) or H_2_O (-H_2_O_2_) and the %GAPDH oxidation (bar graph). **(C-D)** Representative immunoblot analysis of GAPDH oxidation as assessed by mPEG-mal labeling of HEK293 cells with silenced (siSOD1) or unsilenced (siCTRL) *SOD1* **(C)** and % GAPDH oxidation from multiple trials **(D)**. Data represents the average ± SD from five or six biological replicates. **(E-F)** Representative immunoblot analysis of GAPDH oxidation as assessed by mPEG-mal labeling of MCF-7 cells **(C)** and % GAPDH oxidation from multiple trials compared to HEK293 cells **(D)**. Data represents the average ± SD from two and six biological replicates. The statistical significance relative to siCTRL or HEK293 cells is indicated by asterisks using two-tailed Student’s t-test for pairwise comparison. *p<0.05, **p<0.01, ***p>0.001, ****p<0.0001, n.s.= not significant.

**Figure S5.**
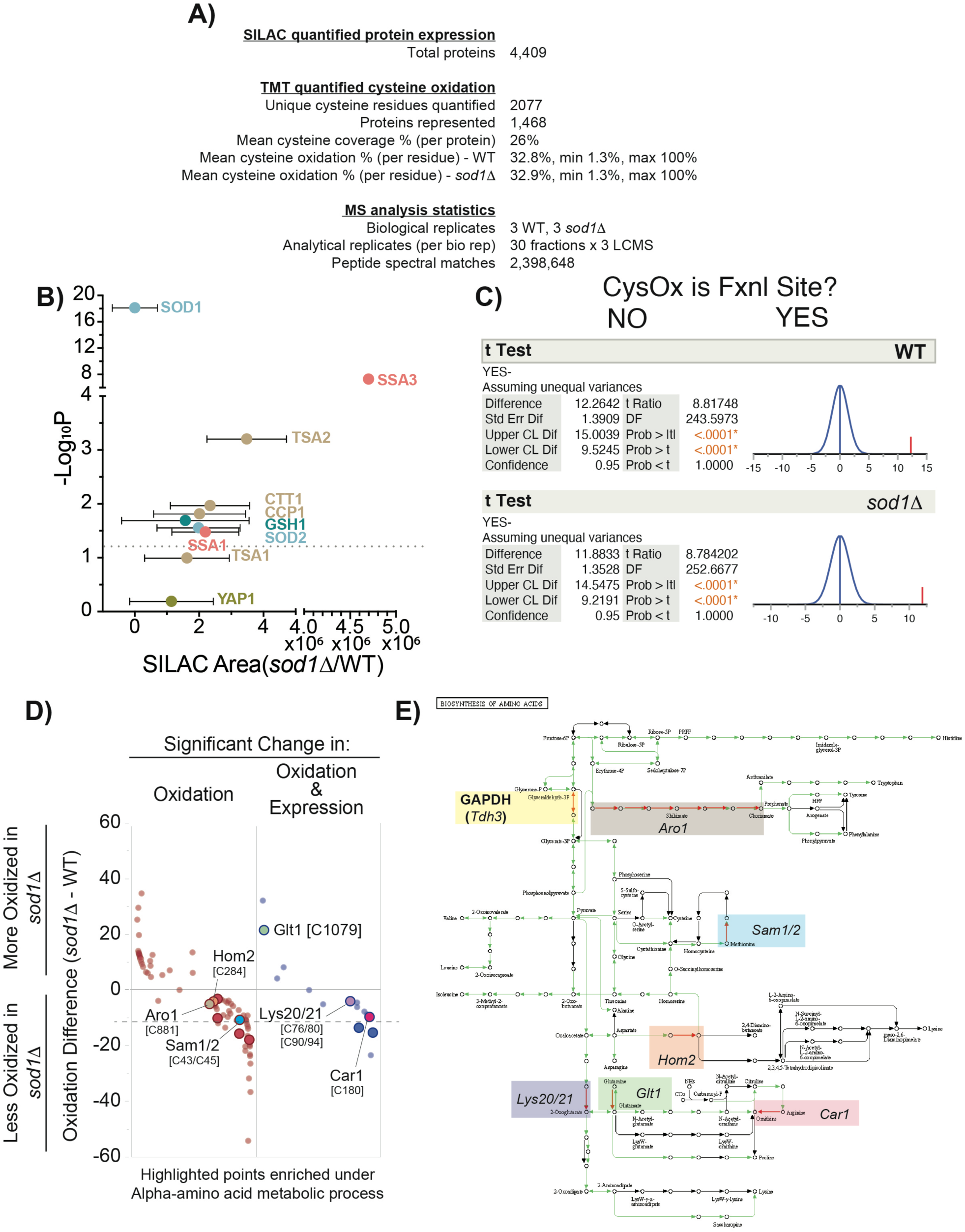
Summary statistics of the redox proteomics experiment and additional analysis of Sod1-dependent protein abundance and oxidation; related to Figure 6. **(A)** Overview statistics for the redox proteomics experiment. **(B)** Subset of data from Figure 6C, showing expression change for antioxidant proteins associated with cellular ROS detoxification. The proteins above the dotted line are significantly more expressed in *sod1*Δ cells. SOD1, Cu/Zn superoxide dismutase; SSA3, Heat Shock protein 73; TSA2, thioredoxin peroxidase; CTT1, cytosolic catalase, CCP1, mitochondrial peroxidase; GSH1, Glutathione synthase; SOD2, Mn superoxide dismutase; SSA1, Heat Shock Protein 71; TSA1, thioredoxin peroxidase. **(C)** Statistical comparison of % oxidation values observed between sites linked with protein function or not. The distributions of cysteine oxidation levels observed for sites required for binding or catalytic activity (YES) versus those that are not (NO). The means of each distribution were compared statistically using a t-test and indicate significantly elevated oxidation for binding and activity-associated residues. **(D** and **E)** Enzymes necessary for amino acid biosynthesis/metabolism undergo significant decrease in oxidation in the absence of SOD1. **(D)** Distribution of significantly changing SOD1-dependent cysteine oxidation sites highlighting proteins (and [cysteine oxidation sites]) that share pathway membership with GAPDH in amino acid biosynthesis processes. Dashed line indicates the average oxidation change observed for GAPDH (Tdh3) C-150 in *sod1*Δ versus WT cells. **(E)** KEGG pathway for Biosynthesis of Amino Acids highlighting enzymes undergoing significant oxidation state change in the absence of *SOD1*.

**Figure S6.**
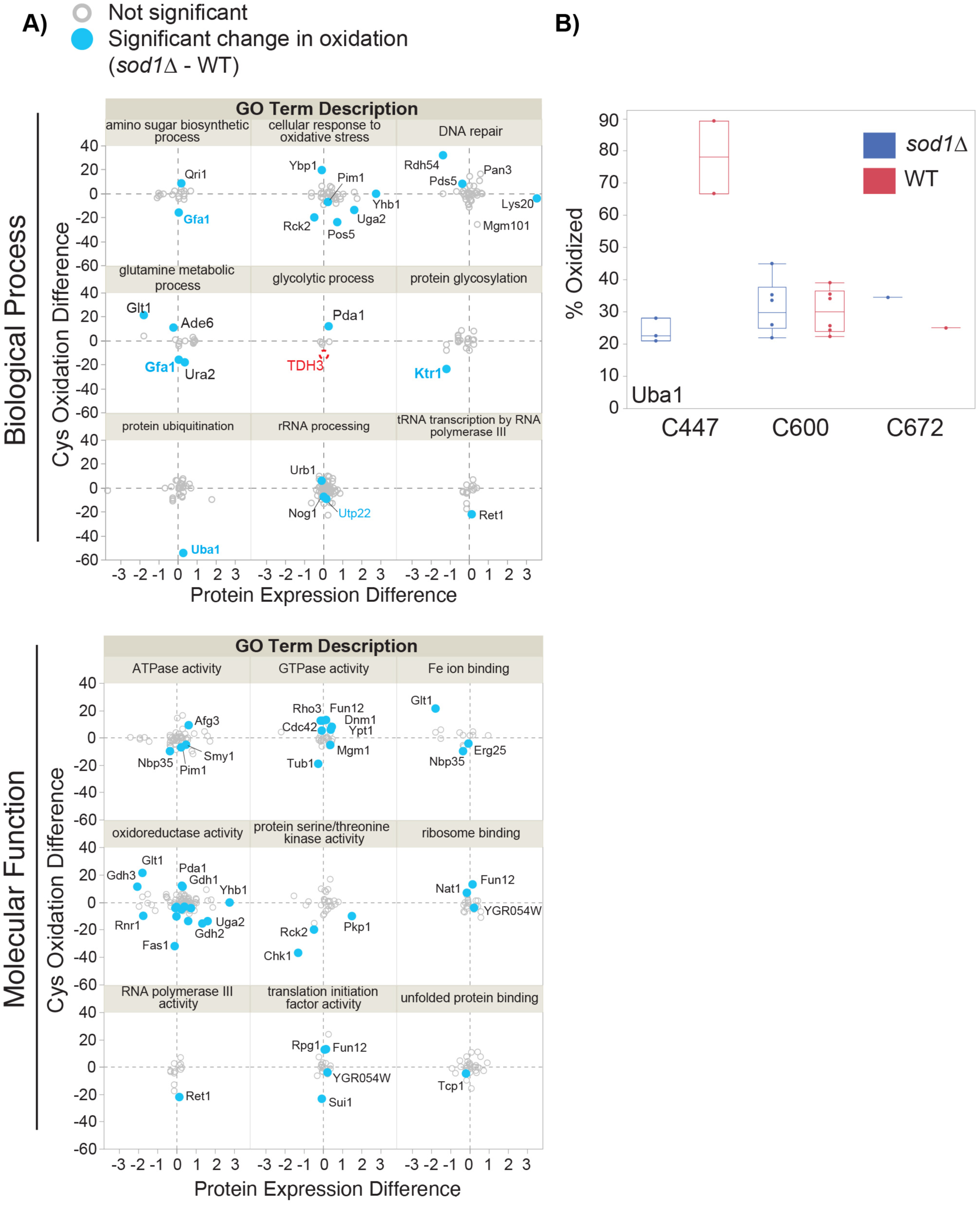
Sod1-dependent protein oxidation organized in terms of molecular function and biological process; related to Figure 6. **(A)** Sod1-sensitive versus insensitive cysteine oxidation sites relative to biological process and molecular function. Plot of cysteine oxidation difference versus protein expression difference for proteins identified in Figure 6F as being significant outliers for their respective GO biological process. For reference, Tdh3-C154 (GAPDH), which was detected but did not release iodoTMT reporters necessary for quantitation is shown based on an average of the oxidation range described earlier for this cysteine (red dashed circle). Protein labels are indicated for those in which a cysteine undergoes significant change between strains. **(B)** Oxidation of Uba1 C-447 is extremely sensitive to SOD1 loss. Replicate measurements for oxidation of all detected cysteine residues in Uba1 are shown. Note that cysteine oxidation differences used in the analyses for Figure 6J are based on differences observed within-replicate, whereas this plot shows all data across all replicates.

**Figure S7.**
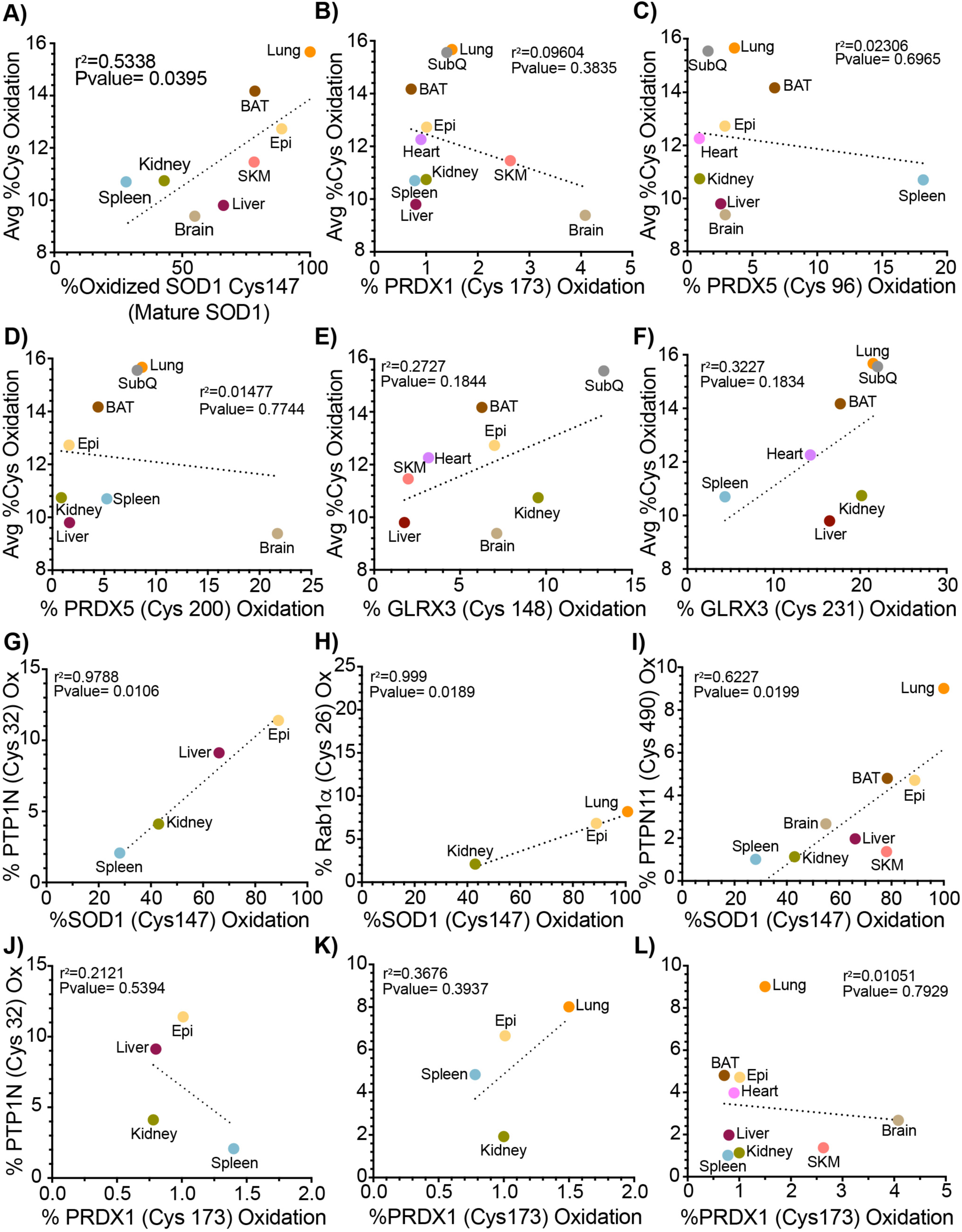
Sod1 maturation correlates with overall and specific protein percentage cysteine oxidation in mice tissues quantified in the Oximouse study. **(A - F)** The correlation between the average of percentage cysteine oxidation (Y axis) and Sod1 Cys147 **(A)**, Peroxiredoxin 1 (PRDX1) Cys173 **(B)**, Peroxiredoxin 5 (PRDX5) Cys96 **(C)**, Peroxiredoxin 5 (PRDX5) Cys200 **(D)**, Glutaredoxin 3 (GLRX3) Cys148 **(E)** and Glutaredoxin 3 (GLRX3) Cys231 **(F)** percentage cysteine oxidation (X axis) form individual tissues is represented as an XY graph. A linear regression analysis gives a coefficient of determination (r^2^) of .5538, 0.09604, 0.02306, 0.01477, 0.02727 and 0.3227, respectively, and a p-value of 0.0395, 0.3835,0.6965, 0.7744, 0.1844 and 0.1834, respectively. **(G - L)** Sod1 Cys147 oxidation correlates with oxidation of PTP1N (Tyrosine-protein phosphatase non-receptor type I) Cys32, mouse homolog of PTP1B (protein tyrosine phosphatase 1B) **(G)**, Rab1*α* Cys26 **(H)** and PTPN11 (Tyrosine-protein phosphatase non-receptor type 11) Cys490 **(I)**. **(J** to **L)** PRDX1 Cys173 oxidation does not correlate with oxidation of PTP1N Cys32, mouse homolog of PTP1B **(J)**, Rab1*α* Cys26 **(K)** and PTPN11 Cys490 **(L).** A linear regression analysis gives a coefficient of determination (r^2^) of 0.9788, 0.0999, 0.6227, 0.2121, 0.3676 and 0.01051, respectively, and a p-value of 0.0106, 0.0189, 0.0199, 0.5394, 0.3937 and 0.7929, respectively. Tissue identity is indicated using distinctive colors and labels. Lung (orange), BAT (brown adipose tissue, brown), Epi (epididymal fat, yellow), SKM (skeletal muscle, coral), liver (dark red), heart (pink), spleen (light blue), kidney (dark green), brain (gold), SubQ (subcutaneous fat, gray). The statistical significance is calculated using linear regression analysis. The Pearson’s correlation coefficient (r) and p-value for each correlation are described in each graph, where p<0.05 corresponds to a significant correlation.

**Table S1.**
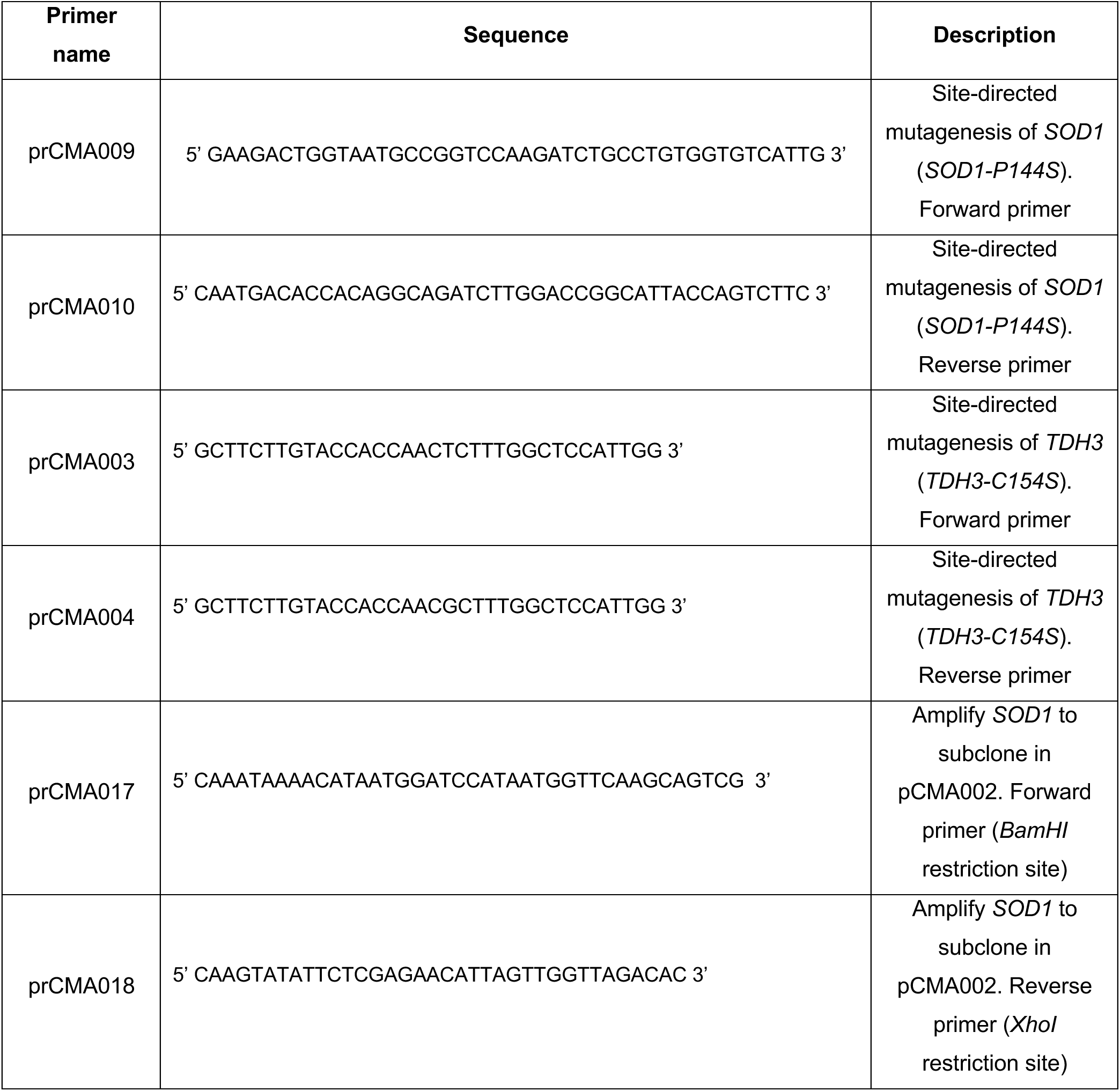

**Table S2: Supplemental MS and gene ontology analysis tables**

## References

1. A. R. Reddi, V. C. Culotta, Regulation of manganese antioxidants by nutrient sensing pathways in Saccharomyces cerevisiae. Genetics 189, 1261–1270 (2011).

2. H. D. Teixeira, R. I. Schumacher, R. Meneghini, Lower intracellular hydrogen peroxide levels in cells overexpressing CuZn-superoxide dismutase. Proceedings of the National Academy of Sciences 95, 7872–7875 (1998).

3. D. H. Flint, j. F. Tuminello, M. H. Emptage, The inactivation of Fe-S cluster containing hydro-lyases by superoxide. J. Biol. Chem. 268, 22369–22376 (1993).

4. S. I. Liochev, I. Fridovich, Superoxide and iron: partners in crime. IUBMB Life 48, 157–161 (1999).

5. J. A. Imlay, Pathways of oxidative damage. Annual Reviews in Microbiology 57, 395–418 (2003).

6. Y. Wang, R. Branicky, A. Noe, S. Hekimi, Superoxide dismutases: Dual roles in controlling ROS damage and regulating ROS signaling. J Cell Biol 217, 1915–1928 (2018).

7. T. Bilinski, Z. Krawiec, L. Liczmanski, J. Litwinska., Is hydroxyl radical generated by the fenton reaction in vivo? Biochem. Biophys. Res. Comm. 130, 533–539 (1985).

8. S. Elchuri et al., CuZnSOD deficiency leads to persistent and widespread oxidative damage and hepatocarcinogenesis later in life. Oncogene 24, 367–380 (2005).

9. E. Chang, D. Kosman, O_2_-dependent methionine auxotrophy in Cu,Zn Superoxide dismutase deficient mutants of *Saccharomyces cerevisiae*. J. Bacteriol. 172, 1840–1845 (1990).

10. E. Gralla, J. S. Valentine, Null mutants of *Saccharomyces cerevisiae* Cu,Zn superoxide dismutase: characterization and spontaneous mutation rates. J. Bacteriol. 173, 5918–5920 (1991).

11. L. A. Sturtz, V. C. Culotta, Superoxide dismutase null mutants of baker’s yeast, Saccharomyces cerevisiae. Methods Enzymol. 349, 167–172 (2002).

12. S. I. Liochev, Commentary: The Role of Iron-Sulfur Clusters in In Vivo Hydroxyl Radical Production. Free radical research 25, 369–384 (1996).

13. S. I. Liochev, I. Fridovich, The role of O2.− in the production of HO.: in vitro and in vivo. Free Radical Biology and Medicine 16, 29–33 (1994).

14. L. B. Corson, J. Folmer, J. S. Strain, V. C. Culotta, D. W. Cleveland, Oxidative stress and iron are implicated in fragmenting vacuoles of *Saccharomyces cerevisiae* lacking Cu,Zn Superoxide dismtuase. J. Biol. Chem. 274, 27590–27596 (1999).

15. C. Montllor-Albalate et al., Extra-mitochondrial Cu/Zn superoxide dismutase (Sod1) is dispensable for protection against oxidative stress but mediates peroxide signaling in Saccharomyces cerevisiae. Redox biology 21, 101064 (2019).

16. A. R. Reddi, V. C. Culotta, SOD1 integrates signals from oxygen and glucose to repress respiration. Cell 152, 224–235 (2013).

17. L. B. Poole, A. Hall, K. J. Nelson, Overview of peroxiredoxins in oxidant defense and redox regulation. Current protocols in toxicology 49, 7.9. 1–7.9. 15 (2011).

18. H. N. Kirkman, G. F. Gaetani, Catalase: a tetrameric enzyme with four tightly bound molecules of NADPH. Proceedings of the national academy of sciences 81, 4343–4347 (1984).

19. H. N. Kirkman, S. Galiano, G. Gaetani, The function of catalase-bound NADPH. Journal of Biological Chemistry 262, 660–666 (1987).

20. S. Sehati et al., Metabolic alterations in yeast lacking copper-zinc superoxide dismutase. Free Radic. Biol. Med. 50, 1591–1598 (2011).

21. K. H. Slekar, D. Kosman, V. C. Culotta, The yeast copper/zinc superoxide dismutase and the pentose phosphate pathway play overlapping roles in oxidative stress protection. J. Biol. Chem. 271, 28831–28836 (1996).

22. S.-M. Jeon, Regulation and function of AMPK in physiology and diseases. Experimental & molecular medicine 48, e245–e245 (2016).

23. D. Peralta et al., A proton relay enhances H2O2sensitivity of GAPDH to facilitate metabolic adaptation. Nature Chemical Biology 11, 156–163 (2015).

24. J. van der Reest, S. Lilla, L. Zheng, S. Zanivan, E. Gottlieb, Proteome-wide analysis of cysteine oxidation reveals metabolic sensitivity to redox stress. Nature communications 9, 1–16 (2018).

25. D. Anastasiou et al., Inhibition of pyruvate kinase M2 by reactive oxygen species contributes to cellular antioxidant responses. Science 334, 1278–1283 (2011).

26. A. R. Mitchell et al., Redox regulation of pyruvate kinase M2 by cysteine oxidation and S-nitrosation. Biochemical Journal 475, 3275–3291 (2018).

27. R. Kunjithapatham, S. Ganapathy-Kanniappan, GAPDH with NAD+-binding site mutation competitively inhibits the wild-type and affects glucose metabolism in cancer. Biochimica et Biophysica Acta (BBA)-General Subjects 1862, 2555–2563 (2018).

28. J. A. Diderich, L. M. Raamsdonk, A. L. Kruckeberg, J. A. Berden, K. Van Dam, Physiological properties of Saccharomyces cerevisiae from which hexokinase II has been deleted. Appl. Environ. Microbiol. 67, 1587–1593 (2001).

29. L. B. Bockus et al., Cardiac insulin signaling regulates glycolysis through phosphofructokinase 2 content and activity. Journal of the American Heart Association 6, e007159 (2017).

30. A. Krüger et al., The Pentose Phosphate Pathway Is a Metabolic Redox Sensor and Regulates Transcription During the Antioxidant Response. Antioxidants & Redox Signaling 15, 311–324 (2011).

31. M. Ralser et al., Dynamic rerouting of the carbohydrate flux is key to counteracting oxidative stress. Journal of biology 6, 10 (2007).

32. M. Ralser et al. (2009) Metabolic reconfiguration precedes transcriptional regulation in the antioxidant response. pp 604–605.

33. A. Kuehne et al., Acute activation of oxidative pentose phosphate pathway as first-line response to oxidative stress in human skin cells. Molecular cell 59, 359–371 (2015).

34. E. Mullarky, L. C. Cantley, “Diverting glycolysis to combat oxidative stress” in Innovative medicine. (Springer, Tokyo, 2015), pp. 3-23.

35. D. Christodoulou et al., Reserve flux capacity in the pentose phosphate pathway enables Escherichia coli’s rapid response to oxidative stress. Cell systems 6, 569–578. e567 (2018).

36. A. A. Shestov et al., Quantitative determinants of aerobic glycolysis identify flux through the enzyme GAPDH as a limiting step. elife 3, e03342 (2014).

37. M. V. Liberti et al., A predictive model for selective targeting of the Warburg effect through GAPDH inhibition with a natural product. Cell metabolism 26, 648–659. e648 (2017).

38. C. C. Winterbourn, M. B. Hampton (2008) Thiol chemistry and specificity in redox signaling.

39. J. C. Juarez et al., Superoxide dismutase 1 (SOD1) is essential for H2O2-mediated oxidation and inactivation of phosphatases in growth factor signaling. Proc. Natl. Acad. Sci. U. S. A. 105, 7147–7152 (2008).

40. P. Jouhten et al., Oxygen dependence of metabolic fluxes and energy generation of Saccharomyces cerevisiae CEN. PK113–1A. BMC systems biology 2, 60 (2008).

41. R. J. Viator et al., Hypoxia-induced increases in glucose uptake do not cause oxidative injury or advanced glycation end-product (AGE) formation in vascular endothelial cells. Physiological reports 3, e12460 (2015).

42. A. Ouiddir, C. Planès, I. Fernandes, A. VanHesse, C. Clerici, Hypoxia upregulates activity and expression of the glucose transporter GLUT1 in alveolar epithelial cells. American journal of respiratory cell and molecular biology 21, 710–718 (1999).

43. B. A. Bruckner, C. V. Ammini, M. P. Otal, M. K. Raizada, P. W. Stacpoole, Regulation of brain glucose transporters by glucose and oxygen deprivation. Metabolism 48, 422–431 (1999).

44. M. Tarsio, H. Zheng, A. M. Smardon, G. A. Martínez-Muñoz, P. M. Kane, Consequences of loss of Vph1 protein-containing vacuolar ATPases (V-ATPases) for overall cellular pH homeostasis. Journal of Biological Chemistry 286, 28089–28096 (2011).

45. J. A. Baron, J. S. Chen, V. C. Culotta, Cu/Zn superoxide dismutase and the proton ATPase Pma1p of Saccharomyces cerevisiae. Biochemical and biophysical research communications 462, 251–256 (2015).

46. D. A. Hanna et al., Heme dynamics and trafficking factors revealed by genetically encoded fluorescent heme sensors. Proc Natl Acad Sci U S A 113, 7539–7544 (2016).

47. L. A. van Leeuwen, E. C. Hinchy, M. P. Murphy, E. L. Robb, H. M. Cochemé, Click-PEGylation–a mobility shift approach to assess the redox state of cysteines in candidate proteins. Free Radical Biology and Medicine 108, 374–382 (2017).

48. L. K. Wood, D. J. Thiele, Transcriptional activation in yeast in response to copper deficiency involves copper-zinc superoxide dismutase. Journal of Biological Chemistry 284, 404–413 (2009).

49. M. Rinnerthaler et al., Yno1p/Aim14p, a NADPH-oxidase ortholog, controls extramitochondrial reactive oxygen species generation, apoptosis, and actin cable formation in yeast. Proceedings of the National Academy of Sciences 109, 8658–8663 (2012).

50. N. M. Brown, A. S. Torres, P. E. Doan, T. V. O’Halloran, Oxygen and the copper chaperone CCS regulate posttranslational activation of Cu,Zn superoxide dismutase. Proc. Natl. Acad. Sci. U. S. A. 101, 5518–5523 (2004).

51. J. M. Leitch et al., Activation of Cu,Zn-superoxide dismutase in the absence of oxygen and the copper chaperone CCS. J. Biol. Chem. 284, 21863–21871 (2009).

52. J. M. Leitch et al., Post-translational modification of Cu/Zn superoxide dismutase under anaerobic conditions. Biochemistry 51, 677–685 (2012).

53. C. White et al., Copper transport into the secretory pathway is regulated by oxygen in macrophages. J. Cell. Sci. 122, 1315–1321 (2009).

54. L. Papa, M. Hahn, E. L. Marsh, B. S. Evans, D. Germain, SOD2 to SOD1 switch in breast cancer. Journal of biological chemistry 289, 5412-5416 (2014).

55. T. P. Dick, M. Ralser, Metabolic remodeling in times of stress: who shoots faster than his shadow? Molecular cell 59, 519–521 (2015).

56. S. Stöcker, M. Maurer, T. Ruppert, T. P. Dick, A role for 2-Cys peroxiredoxins in facilitating cytosolic protein thiol oxidation. Nature chemical biology 14, 148 (2018).

57. Z. A. Wood, L. B. Poole, P. A. Karplus, Peroxiredoxin evolution and the regulation of hydrogen peroxide signaling. Science 300, 650–653 (2003).

58. C. Banks, J. Andersen, Mechanisms of SOD1 regulation by post-translational modifications. Redox biology 26, 101270 (2019).

59. C. J. Banks et al., Acylation of Superoxide Dismutase 1 (SOD1) at K122 Governs SOD1-Mediated Inhibition of Mitochondrial Respiration. Mol Cell Biol 37 (2017).

60. C. K. w. Tsang, Y. Liu, J. Thomas, Y. Zhang, X. F. S. Zheng, Superoxide dismutase 1 acts as a nuclear transcription factor to regulate oxidative stress resistance. Nature communications 5, 3446–3446 (2014).

61. C. K. Tsang et al., SOD1 Phosphorylation by mTORC1 Couples Nutrient Sensing and Redox Regulation. Mol Cell 70, 502–515 e508 (2018).

62. C.-L. Liu et al., Targeting the pentose phosphate pathway increases reactive oxygen species and induces apoptosis in thyroid cancer cells. Molecular and Cellular Endocrinology 499, 110595 (2020).

63. M. L. Gomez, N. Shah, T. C. Kenny, E. C. Jenkins, D. Germain, SOD1 is essential for oncogene-driven mammary tumor formation but dispensable for normal development and proliferation. Oncogene 38, 5751–5765 (2019).

64. R. Somwar et al., Superoxide dismutase 1 (SOD1) is a target for a small molecule identified in a screen for inhibitors of the growth of lung adenocarcinoma cell lines. Proc Natl Acad Sci U S A 108, 16375–16380 (2011).

65. P. Huang, L. Feng, E. A. Oldham, M. J. Keating, W. Plunkett, Superoxide dismutase as a target for the selective killing of cancer cells. Nature 407, 390–395 (2000).

66. J. C. Juarez et al., Copper binding by tetrathiomolybdate attenuates angiogenesis and tumor cell proliferation through the inhibition of superoxide dismutase 1. Clinical cancer research : an official journal of the American Association for Cancer Research 12, 4974–4982 (2006).

67. F. L. Muller, Y. Liu, H. Van Remmen, Complex III releases superoxide to both sides of the inner mitochondrial membrane. J Biol Chem 279, 49064–49073 (2004).

68. D. Han, F. Antunes, R. Canali, D. Rettori, E. Cadenas, Voltage-dependent anion channels control the release of the superoxide anion from mitochondria to cytosol. J Biol Chem 278, 5557–5563 (2003).

69. N. Chandel et al., Mitochondrial reactive oxygen species trigger hypoxia-induced transcription. Proceedings of the National Academy of Sciences 95, 11715–11720 (1998).

70. K. Araki et al., Redox sensitivities of global cellular cysteine residues under reductive and oxidative stress. Journal of proteome research 15, 2548–2559 (2016).

71. U. Topf et al., Quantitative proteomics identifies redox switches for global translation modulation by mitochondrially produced reactive oxygen species. Nature Communications 10.1038/s41467-017-02694-8 (2018).

72. L. I. Leichert et al., Quantifying changes in the thiol redox proteome upon oxidative stress in vivo. Proc Natl Acad Sci U S A 105, 8197–8202 (2008).

73. L. Fu et al., Systematic and quantitative assessment of hydrogen peroxide reactivity with cysteines across human proteomes. Molecular & Cellular Proteomics 16, 1815–1828 (2017).

74. H. Xiao et al., A Quantitative Tissue-Specific Landscape of Protein Redox Regulation during Aging. Cell 180, 968–983.e924 (2020).

75. N. K. Tonks, Protein tyrosine phosphatases: from genes, to function, to disease. Nature reviews Molecular cell biology 7, 833–846 (2006).

76. S.-R. Lee, K.-S. Kwon, S.-R. Kim, S. G. Rhee, Reversible inactivation of protein-tyrosine phosphatase 1B in A431 cells stimulated with epidermal growth factor. Journal of Biological Chemistry 273, 15366–15372 (1998).

77. S. E. Leonard, K. G. Reddie, K. S. Carroll, Mining the thiol proteome for sulfenic acid modifications reveals new targets for oxidation in cells. ACS Chem Biol 4, 783–799 (2009).

78. Y. O. Chernoff, G. P. Newnam, J. Kumar, K. Allen, A. D. Zink, Evidence for a protein mutator in yeast: role of the Hsp70-related chaperone ssb in formation, stability, and toxicity of the [PSI] prion. Mol Cell Biol 19, 8103–8112 (1999).

79. R. D. Gietz, R. H. Schiestl, Applications of high efficiency lithium acetate transformation of intact yeast cells using single-stranded nucleic acids as carrier. Yeast 7, 253–263 (1991).

80. F. Ness et al., Sterol uptake in Saccharomyces cerevisiae heme auxotrophic mutants is affected by ergosterol and oleate but not by palmitoleate or by sterol esterification. Journal of bacteriology 180, 1913–1919 (1998).

81. E. Luk, M. Carroll, M. Baker, V. C. Culotta, Manganese activation of superoxide dismutase 2 in Saccharomyces cerevisiae requires MTM1, a member of the mitochondrial carrier family. Proc. Natl. Acad. Sci. USA 100, 10353–10357 (2003).

82. L. Flohe, F. Otting, “Superoxide dismutase assays” in Methods in enzymology: oxygen radicals in biological systems, L. Packer, Ed. (Academic press, New York, 1984), vol. 105, pp. 93-104.

83. D. S. Perlin, S. L. Harris, D. Seto-Young, J. E. Haber, Defective H(+)-ATPase of hygromycin B-resistant pma1 mutants fromSaccharomyces cerevisiae. J. Biol. Chem. 264, 21857–21864 (1989).

84. D. m. Wessel, U. Flügge, A method for the quantitative recovery of protein in dilute solution in the presence of detergents and lipids. Analytical biochemistry 138, 141–143 (1984).

